# The human branchpoint-interacting stem loop sequence and structure regulates U2 snRNA expression, branchpoint recognition, and transcriptome

**DOI:** 10.1101/2025.08.31.673180

**Authors:** Meredith B. Stevers, Sol Katzman, Melissa S. Jurica

## Abstract

During pre-mRNA splicing, the branch helix forms when U2 snRNP engages with introns to initiate spliceosome assembly. Its formation is mutually exclusive with the branchpoint-interacting stem loop (BSL) present in U2 snRNA. While BSL structure impacts splicing with the strict consensus branchpoint sequence of yeast introns, its influence in the flexible context of human branchpoints is unknown. We employed an orthogonal U2 snRNA and splicing reporter to examine the impact of perturbing BSL base-pairing and found differential effects on both orthogonal U2 snRNA expression and reporter splicing, indicating that BSL structure influences the biogenesis of U2 snRNP and its function in splicing. Furthermore, high complementarity between the branchpoint sequence and U2 snRNA increases splicing efficiency with wildtype and stabilized BSL, but not when BSL base-pairing is reduced. These data are consistent with complementarity between the intron and the loop of the BSL driving intron-mediated unwinding of the BSL stem. Finally, we investigated transcriptome-wide effects of expressing U2 snRNA with either a cancer-associated BSL mutation or with an altered branchpoint recognition sequence. Similar changes in splicing and gene expression suggests that while altered U2 snRNA is tolerated, cells respond by upregulating genes linked to oncogenic pathways.

**Main Conclusions:**

- U2 snRNA BSL base-pairing influences branchpoint sequence recognition and splicing of an orthogonal splicing reporter
- Propose a model for branchpoint sequence complementarity inducing BSL unwinding to promote splicing efficiency
- U2 snRNA BSL and BPRS mutations alter cellular gene expression regulation and splicing

**GRAPHICAL ABSTRACT:** 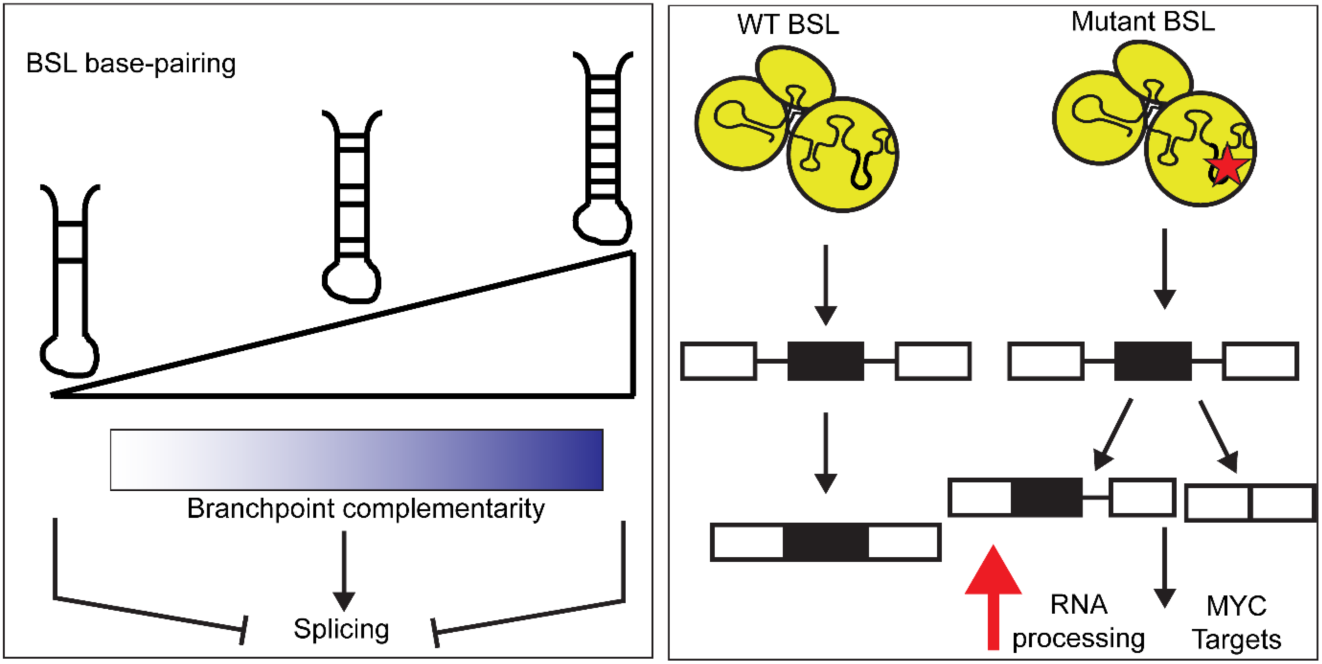

## INTRODUCTION

A critical step of intron recognition by the spliceosome involves the ATP-driven engagement of U2 snRNP with the branchpoint region resulting in an RNA duplex between the U2 snRNA branchpoint recognition sequence (BPRS) and intron branchpoint sequence, termed the branch helix. Formation of the branch helix is critical for splicing because it bulges the branchpoint adenosine necessary for the first chemical step of splicing and indirectly defines the first downstream AG dinucleotide as the intron’s 3′ splice site (1). Base-pairing between U2 snRNA and intron is clearly fundamental to branch helix formation. The nearly universal UACUAAC branchpoint sequence found in *S. cerevisiae* introns perfectly complements the conserved GUAGUA of the U2 snRNA BPRS. Additionally, foundational studies in both yeast and human splicing further demonstrated that compensatory mutations of U2 snRNA suppressed branchpoint sequence changes that disrupted splicing (2–4). Furthermore, the degree of complementarity between U2 snRNA and branchpoint sequence correlates with splicing levels in the context of human minigene splicing assays (5, 6). However, the weaker branchpoint consensus sequence in many other organisms, including the NNyUNAy of human introns, suggests that other factors are involved in intron recognition (7, 8).

The molecular mechanism of branch helix formation and its regulation remains mysterious. Cryo-EM models of U2 snRNP before and after branch helix formation suggest that changes in U2 snRNA structure must also accompany the process. In the standalone human 17S U2 snRNP, the BPRS of U2 snRNA overlaps with the loop of a poorly resolved helix termed the branchpoint-interacting stem loop (BSL) and is surrounded by U2 snRNP proteins, HTATSF1 and DDX46 (9–11). The imperfect stem consists of five Watson-Crick (WC) base pairs interspersed with single G-U, U-U and C-A interactions. In spliceosome models caught after the branch helix has formed, the BSL is absent, and its nucleotides are instead paired with the intron. BSL unwinding could therefore serve as a regulatory step in branch helix formation.

Consistent with this idea, the first studies characterizing the BSL in yeast reported genetic interactions between BSL mutants, Cus2 (HTATSF1 ortholog) and Prp5p (DDX46 ortholog) as evidence of a role for BSL structure and unwinding in the fidelity of branchpoint sequence recognition (12). Intriguingly, a recent report identified a recurrent U2 snRNA mutation (C28U) in cancers that disrupts a WC base pair pointing to the relevance of BSL structure to cell physiology (13). However, questions about the role of the BSL in human splicing remain to be resolved.

To explore the question of does base-pairing between U2 snRNA and the intron have a role in BSL unwinding, and if so, how does the highly variable branchpoint sequence of human introns effect this, we endeavored to determine the effect of perturbing the sequence and base-pairing potential of the human BSL on splicing. Our data indicate a role for BSL dynamics in U2 snRNP biogenesis, and support a model in which complementarity of a 3-nucleotide toehold between the U2 snRNA BPRS and branchpoint sequence controls the initiation of BSL unwinding by the intron.

We also investigated the global effects of perturbing U2 snRNA sequence on splicing and gene expression. Although we do not identify a strong splicing signature, we speculate a mild general splicing defect results in lowered expression of many genes. Cells may try to counteract this defect by upregulating genes important for pre-mRNA processing, translation regulation, and protein folding, which may explain the relevance of U2 snRNA mutations in cancer.

## MATERIAL AND METHODS

### Orthogonal splicing reporter and U2 snRNA expression plasmids

Based on the system developed by Wu and Manley (4, 14), the SV40 Large T antigen gene was amplified from Addgene plasmid #2297 and subcloned into the AflII and NotI sites of Addgene plasmid #44170 downstream of a CMV promoter. To create the branchpoint splicing reporters, the small T branchpoint was mutated by around-the-horn site-directed mutagenesis and verified by Sanger sequencing. For the orthogonal U2 snRNA construct, the RNU2-1 locus including its Pol II promoter and 3’ processing site was amplified from HeLa genomic DNA (15) and subcloned into the BglII site of Addgene plasmid #44170. The orthogonal ACU branchpoint recognition sequence and BSL mutations were introduced by around-the-horn site-directed mutagenesis and verified by Sanger sequencing. For lentiviral transduction, the wildtype and orthogonal RNU2-1 locus was subcloned into the XbaI and NotI sites of a custom pSico lentiviral transfer vector (gift from Susan Carpenter lab) containing an EF1A promoter inducing expression of Zeocin resistance-T2A-GFP. The BSL C28U mutation was introduced into the wildtype U2 snRNA sequence by around-the-horn site-directed mutagenesis and verified by Sanger sequencing. Primer sequences for cloning and mutagenesis are included in Supplementary Table 1.

### Branchpoint sequence analysis

SVM_BPfinder(16) (https://github.com/comprna/SVM-BPfinder) was used to analyze the branchpoint and SVM scores of branchpoints with different BPRS complementarities.

Branchpoint sequences of various complementarity to the orthogonal U2 snRNA were curated and equivalent sequences to the wildtype branchpoint sequence were evaluated at the same location within the context of the small T intron. Program options run with each tested intron include, -s Hsap for homo sapiens, -l 100 for read length of intron, and -d 10 for distance (nt) allowed between branchpoint A and 3’ splice site.

### Transfections and RNA isolation for splicing reporter experiments

HEK293 cells (gift from Doug Kellogg lab) were cultured at 37°C, 5% CO_2_ in 24-well tissue culture plates (Falcon, cat 353047) in Dulbecco’s Modified Eagle Medium (Gibco, cat 12100061) supplemented with 10% FBS (Hyclone, cat SH3054103). Cells at 60-80% confluency were transiently transfected with a 3:1 v/v ratio of Fugene 6 transfection reagent (Promega, cat E2693) to DNA containing 150 ng of splicing reporter plasmid and 350 ng of U2 snRNA expression plasmid. After 72 hours, total RNA was isolated from the transfected cells by Trizol extraction (Invitrogen, cat 15596018) and ethanol precipitation. Transfections were performed with a minimum of three biological replicates.

### Reverse Transcription

Total RNA was treated with RQ1 DNase (Promega, cat M6101) followed by phenol:chloroform:iso-amyl alcohol (25:24:1 v/v) extraction, chloroform:iso-amyl alcohol (24:1 v/v) extraction and ethanol precipitation. The DNase-treated RNA was resuspended in distilled water and quantified by UV absorbance with a NanoDrop 2000 (Thermo Scientific, cat ND-2000). RNA integrity was confirmed before and after DNAse treatment by agarose gel electrophoresis. First-strand cDNA synthesis was performed by first annealing primers specific to SV40 or U2 snRNA (Supplementary Table 1) in a 14 µl reaction containing 800 ng of DNase-treated RNA, 1 µl of 2 µM primers, and1 µl of 10 mM dNTPs at 65°C for 5 mins and placed on ice for 5 mins. Extension buffer containing 4 µl 5X First strand buffer (Invitrogen), 1 µl 0.1 M DTT, and 1 µl of a modified MMLV-RT was added and reactions were incubated at 55°C for 50 mins and 85°C for 5 min. The modified MMLV-RT containing D524G, D583N, and E562Q mutations was expressed and purified in-house, and stored in 20 mM Tris-HCl (pH 7.5), 150 mM NaCl, 0.1 mM EDTA, 1 mM DTT, 0.01% v/v NP-40, 50% v/v glycerol at a working concentration of 1mg/ml.

### U2 snRNA expression primer extension

To quantify orthogonal U2 snRNA expression relative to endogenous, U2 snRNA cDNA was first amplified from 40 ng of first-strand reverse transcription product with primers complementary to the 5’ and 3’ end of U2 snRNA sequence (Supplementary Table 1) by 25 cycles of PCR with Taq polymerase (gift from David Feldheim lab), and isolated with the NucleoSpin Gel and PCR Clean-up kit (Macherey-Nagel, cat 740609.250). A 5’-end-labeled primer was generated by incubating 10 pmol of a DNA oligonucleotide complementary to nucleotides 39-54 with γ-^32^P ATP and PNK4 kinase (Thermo Scientific cat EK0031), followed by size exclusion with a Sephadex G-25 (Sigma-Aldrich, cat G2580-50G) spin column. For primer extension reactions, 50 ng of U2 snRNA amplified cDNA and 0.1 pmol of labeled primer were incubated in 12 µl with annealing buffer (83.3 mM Tris-HCl pH 7.9, 125 mM KCl) at 95°C for 2 minutes, 37°C (33°C for M9, M8/9 primer (Supplementary Figure 4A)) for 10 minutes, and room temperature for 30 minutes, and then supplemented with 8 µl of extension mix to yield a final concentration of 10 mM DTT, 3 mM magnesium chloride, 0.1 mM dATP, dTTP, dCTP, and ddGTP and 2 units of AMV reverse transcriptase (NEB, cat M0277S). The reactions were incubated at 42°C for 70 mins followed by addition of 10 volumes of 0.3 M sodium acetate pH 5.2, 0.5 M EDTA, and 0.05% SDS and 20 ng glycogen. As a negative control, the template and extension mix were substituted by water. As a positive control, the amplified U2 cDNA template was replaced with a correlating amplicon from the orthogonal U2 snRNA expression plasmid.

Extension products were isolated by ethanol precipitation, resuspended in 95% v/v formamide, 20 mM EDTA, 0.01% bromophenol blue and 0.01% cyan blues and separated on a 15% (v/v) polyacrylamide 7 M urea 1X TBE gel that was dried onto Whatman paper and visualized by phosphor imaging. Fiji (Image J) (version 1.53t) was used to determine pixel intensities of the ddGTP extension stop bands at C35 and C28, which were adjusted for incomplete ddGTP incorporation based on C28 intensity caused by C35 readthrough in the positive control. The resulting values were divided by the sum of C35 and C28 band intensities to express the ratio of orthogonal U2 snRNA relative to total U2 snRNA. Statistical analysis of differences in expression was performed in Prism 10 (Graph Pad) using a paired two-way Student’s T-test to determine *p*-values with an alpha = 0.05.

In transduced HEK293T cells, the same strategy was used to assess expression of exogenous U2 snRNA expression, except that the radiolabeled primer for the C28U mutant was complementary to nucleotides 32-47 and ddATP was substituted for ddGTP in the extension mix.

### Quantitative PCR for the orthogonal splicing reporter

TaqMan hydrolysis (Thermo Fisher) probes specific to the splice junctions of large T and small T antigen isoforms, and to exon 2 were synthesized with 5’ FAM and 3’NFQ-MGB quencher (Eurofins) (Supplementary Table 1). Amplicon size and primer specificity for the correct spliced product were both confirmed by gel electrophoresis. Serial dilutions of cDNA (1:5 to 1:80) against primer (4.5-18µM) and probe (2.5-10µM) concentrations were tested to optimize qPCR efficiency to 110-120% with r^2^ between 0.9 and 1. Reactions were prepared by combining 1 µl of 1:10 diluted cDNA to 19 ul of qPCR reaction (5 ul 4X TaqMan Fast Virus 1-Step Master Mix (Applied Biosystems, cat 4444432), with 0.5 ul probe 2.5 uM to 10 uM, and 1 ul of each forward and reverse primers 4.5 uM to 18 uM (Supplementary Table 1) and then incubated for 1 cycle of 20 sec at 95°C followed by 40 cycles of 15 sec at 95°C and 1 min at 60°C with a QuantStudio 6 Pro Real-Time PCR system (Applied Biosystems). Three technical replicates were prepared for each biological sample tested. RT-PCR reactions using DNase-treated RNA as template were used to rule out DNA contamination. Design and Analysis software 2.6.0 (Applied Biosystems) was used to determine Cq values for reactions. Baseline correction was derived from the linear amplification plot and then the log-based amplification plot was used to set threshold values within the linear phase where the technical and biological replicates exhibited the least variability and above the background signal of a no template control reaction. Reactions with Cq values that differ by >0.5 relative to other technical replicates were removed as outliers before calculating an average Cq value for each biological replicate.

A modified ΔΔCt method (17) using exon 2 as the reference control was used to determine the small T and large T reporter splicing efficiency for U2 variants relative to empty vector control for each group of transfections. Specifically, the Cq value for splice junction probes of a sample was subtracted from the Cq value for the corresponding exon 2 probe to yield ΔCt from which the ΔCt value for empty vector control is subtracted to yield a -ΔΔCt value that represent the log2-fold change in splicing efficiency relative to empty vector. Statistical analysis of ΔΔCt value comparisons was performed in Prism 10 (Graph Pad) using paired two-way Student’s T-test to determine *p*-values with an alpha level of 0.05.

### Lentiviral transduction and selection of HEK293T U2 mutant cell line

To generate lentivirus for exogenous U2 snRNA expression, HEK293T cells (gift from Angela Brooks lab) were seeded into 6 well plates (BioLite, cat 130184) at 1×10^6^ cells/well in DMEM supplemented with 10% FBS. After 24 hours, cells were transfected with 0.75 µg psPAX2 and 0.25 µg pMD2.G (lentiviral packaging plasmids gifted from Joseph Costello lab), and 1 ug of custom pSico lentiviral transfer vector containing BPRS or BSL mutant U2 snRNA expression locus using X-tremeGENE HP (Sigma-Aldrich, cat 06366244001) transfection reagent in Opti-MEM (Gibco, cat 31985062) according to manufacturer directions. After a 24-hour incubation, media was replaced with DMEM /10% FBS. Virons were collected 72 hours post-transfection by filtering the media through 0.2 µM PES (Celltreat, cat 229746), aliquoted and stored at -80°C.

Replicate mutant U2 snRNA cell lines were generated by first seeding HEK293T cells in 12 well plates (CELLSTAR, cat 665180) at 1.5×10^5^cells/well in DMEM/10% FBS. After 24 hours, threefold serial dilutions of stock virus and 4 µg/ml polybrene were added for each U2 snRNA mutant construct and empty vector control line, and the cells incubated for 24 more hours. The media was then replaced with fresh DMEM/10% FBS and the cells grown for 72 hours. Transduced cells were selected by growth and maintained in media supplemented with 300 µg/ml Zeocin (gift from Susan Carpenter lab) (Thermo Fisher, cat R25001). For each replicate, the polyclonal cell line from viral dilutions yielding the highest GFP fluorescent signal were selected for further culture, characterization and downstream RNA-seq analysis.

### Short-Read RNA-seq

For each genotype transduced cell line, total RNA was extracted with Trizol (Invitrogen, cat 15596018) and ethanol precipitation (n=2 for one replicate polyclonal cell line and n=1 for the other replicate). RNA concentration and RIN numbers (ranging from 7.1 to 9.2) were determined by Agilent 2100 Bioanalyzer (Agilent Technologies). UC Davis DNA Technologies and Expression Analysis Core Laboratory prepared poly-A strand specific libraries and sequenced 58 million paired end read pairs by NovaSeq S4 (PE150).

### RNA-seq Data Analysis

Fragment quality was initially checked using Salmon (v1.10.3) (18) by quasi-mapping reads to an index built from the human reference genome release 45 primary comprehensive gene annotation (GRCh38) from Gencode (https://www.gencodegenes.org). The Salmon output was converted into HTML format by Multiqc (v1.27) (19) and reads were confirmed to distribute around ∼ 400 bp. Raw fastq read quality was assessed using FastQC (v0.12.1) (20) and remaining Illumina adaptors and low-quality reads were detected and removed by trim_galore (v0.6.10) (21). Trimmed reads were then aligned to Gencode human reference genome release 47 primary comprehensive gene annotation (GRCh38) using STAR (v2.7.11b) (22) with the following options: --outFilterMultimapNmax 20, --alignIntronMax 1000000, --alignMatesGapMax 1000000, --alignSJDBoverhangMin 1, --limitSjdbInsertNsj 2000000, --outSAMattributes NH HI AS nM jM, --alignIntronMin 20, --outSAMtype BAM SortedByCoordinate, --twopassMode Basic, and --quantMode TranscriptomeSAM GeneCounts. Bam files were indexed using samtools (v1.21 with htslib 1.21) (23) and bigwig files were created for the forward and reverse reads using deeptools (v3.5.6) (24). Read counts were normalized using the inverse of the DESeq2 size factors.

### Differential Expression Analysis

Differential gene expression analysis was performed using DESeq2 (v1.42.1) (25) after extracting reverse stranded read counts from the STAR alignment and removing genes with no read counts across all samples. Reference condition was set to the empty-vector control cell line samples. Results were FDR adjusted with alpha = 0.05. Significantly upregulated or downregulated genes were evaluated for GO term enrichment using Enrichr (26–28).

### Differential Splicing Analysis

Novel splice sites were incorporated into the v47 GTF file using Stringtie (v3.0.1) (29) from the BAM flies. Splice sites that did not have at least 3 bases on either side were filtered out with –a 3. GTFs were created for each sample and merged. Splicing analysis was then performed using junctionCounts (v1.0.0) (30) with the merged GTF file using default settings except for DEXSeq where --min_jc 10, --min_psi 0.1, and --ri_span 0.1. Significant splice events were extracted and analyzed further using bam coverage tracks on the UCSC genome browser and bam files loaded in IGV (v 2.17.4) (31). Sashimi plots of analyzed events were generated with IGV.

## RESULTS

### Mutations within the BSL affects both snRNA expression levels and reporter splicing in an orthogonal-U2 system

Modifying U2 snRNA in human cells is challenging. U2 snRNA is encoded by multiple sequence variants, with the predominant variant RNU2-1 transcribed from a multi-copy gene array, precluding simple genetic alteration (13, 32–35). Therefore, we adopted an orthogonal U2 splicing system developed by Wu and Manley employing a modified SV40 Large T antigen gene with two alternative splice isoforms termed large T and small T (4, 14). To recreate the orthogonal splicing system, we constructed RNU2-1 expression plasmids with a BPRS containing either the normal UAG (U2-WT) or a ACU transversion (U2-Ortho) along with splicing reporter plasmids containing the SV40 Large T antigen gene in which the branchpoint required for small T intron splicing is complementary to the normal or orthogonal U2 (BP-WT and BP-Ortho, Figure 1A and B). Splicing of the BP-Ortho small T isoform therefore reports on the functionality of the exogenous U2-Ortho snRNA. To confirm the functionality and specificity of the orthogonal splicing system, we transfected HEK293 cells with different combinations of these plasmids and assessed both U2-Ortho expression and reporter splicing. Primer extension with cDNA of U2 snRNA as template and ddGTP produced an extension stop mapping to the orthogonal nucleotide C35 only with samples from U2-Ortho transfected cells, while all samples exhibited a stop at nucleotide C28 present in all U2 snRNA versions (Figure 1C, compare lanes 3 and 4 with 1 and 2). Based on the ratio of the C35 to C28 band intensities, we estimate that 15% of cellular U2 snRNA contained the orthogonal BPRS. RT-qPCR with probes specific to the large and small T splice products demonstrated that splicing of the BP-Ortho reporter is significantly higher with U2-Ortho transfection relative to U2-WT (Figure 1D, compare 2 and 4). In contrast, splicing of the BP-WT reporter occurs regardless of the identity of the U2 expression construct transfected (Figure 1D, compare lanes 1 and 3), likely due to the contribution of endogenous U2 snRNA. From these results, we conclude that U2-Ortho is expressed at levels sufficient to support orthogonal reporter splicing and that efficient splicing of the BP-Ortho reporter relies on the presence of U2-Ortho. Furthermore, large T splicing levels were nearly equivalent across all transfections, indicating that expression levels of SV40 large T antigen were stable and did not contribute to the changes observed in small T splicing (Supplementary Figures 1, 3 and 4).

**Figure 1.**
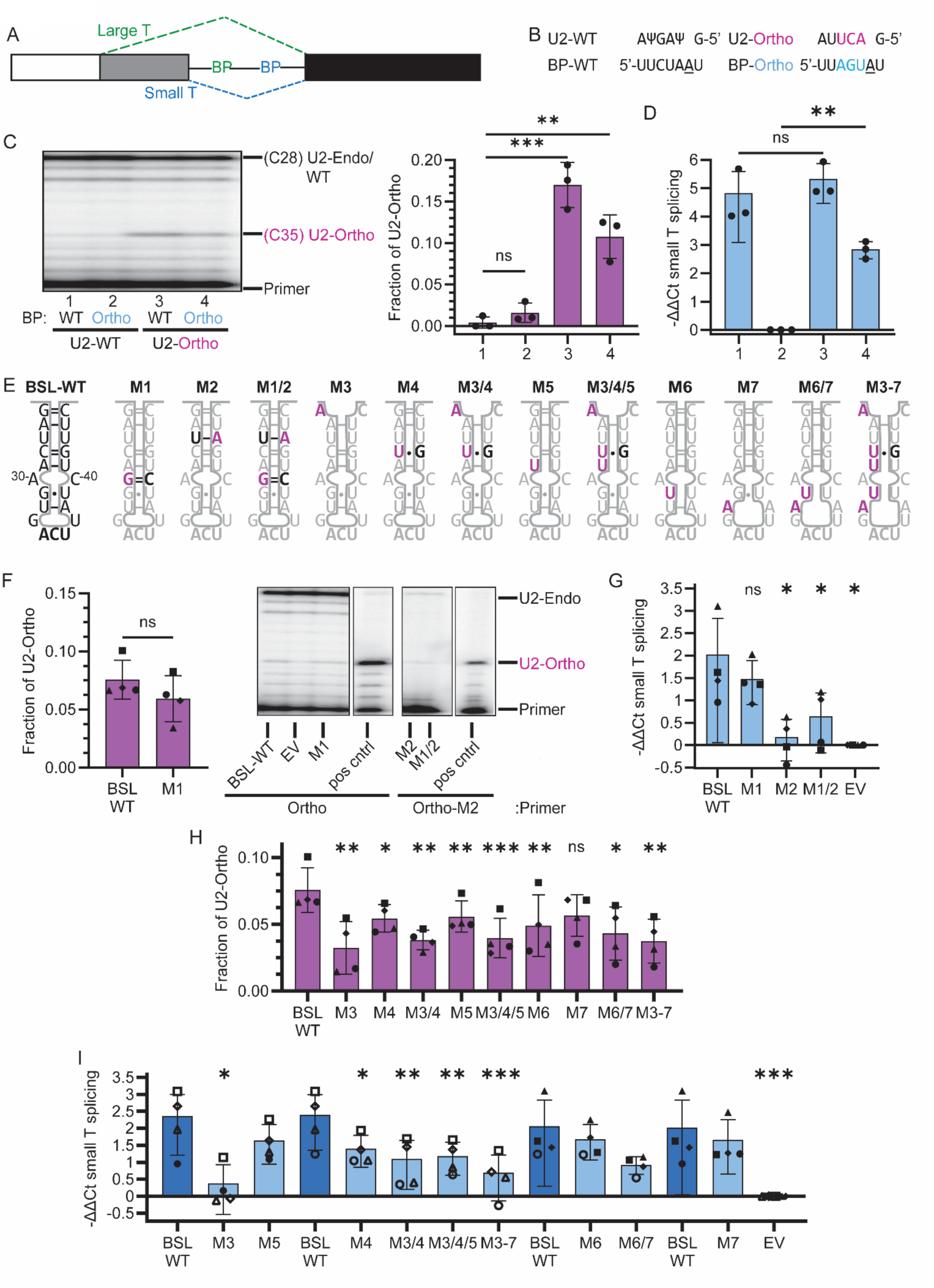
Perturbing BSL base-pairing potential decreases Ortho U2 snRNA stable expression and reporter splicing. (**A**) Simplified schematic of SV40 T antigen alternative splicing. Splicing for the large T isoform is highlighted by the green dashed line and the small T isoform and branchpoint is in blue. (**B**) BPRS and branchpoint sequences of wildtype and orthogonal splicing. U2 snRNA wildtype (U2-WT) and orthogonal (U2-Ortho) are shown above the branchpoint sequences of the SV40 small T wildtype (BP-WT) and orthogonal (BP-Ortho) introns. Orthogonal nucleotides are highlighted in pink and blue. The branchpoint adenosine for both is underlined. (**C**) Expression of exogenous U2 snRNAs in HEK293 cells. Left: Representative PAGE of radiolabeled U2 snRNA primer extension analysis from cells transfected with U2-WT and U2-Ortho. Extension stops for endogenous/wildtype U2 snRNA and U2-Ortho are labeled. Right: Fraction of U2-Ortho relative to total U2 snRNA quantified from primer extension stops; n=3. (**D**) Difference in small T intron splicing by RT-qPCR for samples shown in (C). -ΔΔCt is derived from small T splice junction probe normalized to exon2 and relative to sample 2 (BP-Ortho and U2-WT). (**E**) Schematic of orthogonal human U2 snRNA BSL wildtype and mutant sequences with predicted base-pairing. Mutations are indicated in pink. (**F**) Expression levels of BSL stabilizing U2-Ortho mutants. Fraction of U2-Ortho relative to total U2 snRNA was determined as described in (C); n=4. (**G**) Difference in small T intron splicing by RT-qPCR for samples shown in (F). -ΔΔCt is derived from small T splice junction probe normalized to exon2 and relative to empty vector (EV). (**H**) Expression levels of BSL destabilizing mutant U2-Ortho; n=4. (**I**) Difference in small T intron splicing by RT-qPCR for samples shown in (H). - ΔΔCt is derived as in (G). Lanes are grouped by transfection batch for comparison. Matching closed and open shapes are technical replicates of one biological replicate. In all cases, error bars represent standard deviation.

We next set out to test the hypothesis that the base-pairing potential of the BSL influences BSL dynamics to affect branchpoint recognition and splicing by introducing mutations in the sequence of U2-Ortho to either increase (M1 to M1/2) or decrease the base-pairing interactions within the BSL stem (M3 to M3-7) (Figure 1E). The mutant U2-Ortho expression plasmids were each transfected into cells along with the BP-Ortho splicing reporter and total RNA analyzed for U2 snRNA expression and reporter splicing. U2-Ortho with a wild-type BSL stem (BSL-WT) comprised ∼7.5% of cellular U2 snRNA and consistently yielded a log2-fold increase of ≥2 for reporter splicing relative to empty vector (Figure 1F-I). M1 (A30G), which changes the A-C bulge between nucleotides 30 and 40 to a stabilizing G-C base pair, was expressed at slightly reduced levels and supported reporter splicing that is not significantly different from BSL-WT (Figure 1F and G). M2 (U43A) converted a U-U interaction between nucleotides 27 and 43 to a Watson-Crick base pair and resulted in low U2-Ortho expression with a correlating decrease in reporter splicing. The M1/2 combination of both mutations also displayed low U2-Ortho expression and splicing levels. We conclude that increased base-pairing at some positions of the BSL likely interferes with splicing due to increased turn-over of the altered U2 snRNA, hypothetically because BSL structural dynamics are involved in the incorporation or maintenance of U2 snRNA into the snRNP. However, some perturbation to the BSL is tolerated relative to both U2 snRNP assembly and function because M1 is both expressed and confers reporter splicing.

We next tested a series of mutations aimed at decreasing the base-pairing potential of the BSL. Three single mutations were introduced within the base of the stem (M3 G25A, M4 C28U, M5 A29U) and two were adjacent to the loop of the stem (M6 G31U, M7 U32A) (Figure 1E). We also tested combinations M3/4, M6/7 and M3/4/5, and the quintuple M3-7. The changes generally lowered U2-Ortho expression, but to different degrees (Figure 1H). Introduction of M4 or M5 only partially reduced U2-Ortho expression, while introduction of M3 had a more significant effect. The effect of M3 appears to be dominant because the combinations with M4 and M5 (M3/4 and M3/4/5) resulted in similar lower U2-Ortho levels. Mutations near the loop of the stem had less of an effect on U2-Ortho levels, with M6 and M7 resulting in a small or non-significant decrease in expression. In combination, however, the M6/7 mutations resulted in a decrease of U2-Ortho. The expected full destabilization of the BSL with M3-7 roughly halved the level of U2-Ortho relative to BSL-WT. These results indicate that loss of most single base-pairing interactions within the BSL does not severely impact steady-state levels of the U2 snRNA. Still, the reduction of U2-Ortho with M3 and M3-7 indicates that certain changes to the BSL structure can negatively affect incorporation or maintenance of U2 snRNA in the snRNP.

The BSL-destabilizing mutations differentially affected splicing of the BP-Ortho reporter (Figure 1I). M6 and M7 on their own had no significant effect. The M6/M7 combination results in decreased reporter splicing, but the impact of BSL structure on splicing cannot be disambiguated from the decrease on U2-Ortho expression. Similarly, decreased reporter splicing of M5, M3/4, and M3/4/5 also correlated with lower U2-Ortho expression. With M3 and M3-7, however, we detect no BP-Ortho reporter splicing above background. This loss of splicing could be due to low U2-Ortho expression, but this explanation is likely insufficient considering that other mutants with comparable expression levels support reporter splicing. The M3 and M3-7 mutations may therefore also interfere with the function of U2 snRNA in splicing. In the cryo-EM structures of U2 snRNP, a tryptophan extending from SF3A3 stacks on the base pair between G25 and C45 and appears to constrain the location of the base of the BSL stem (9–11). Disruption of this base pair with M3 may therefore affect either BSL positioning or SF3A3’s assembly into U2 snRNP. Based on the relatively small effect of M4 on U2-Ortho expression and more significant decrease in reporter splicing, this mutation also appears to interfere with U2 snRNA function. We note that the same C28U change was recently identified in U2 snRNA in pancreatic, prostate and several hematological cancers (13).

### Branchpoint complementarity regulates splicing efficiency of orthogonal U2 snRNA

Complementarity between the U2 BPRS and branchpoint sequence of an intron is critical in yeast, but less constrained for splicing in humans (7, 8). In yeast, BSL mutations genetically interact with branchpoint sequence changes to suggest a role for the BSL in branchpoint sequence recognition (12). To test whether a similar relationship holds in the human system, we tested nucleotide changes within four of the six nucleotides flanking the branchpoint adenosine (Figure 2A). In all cases, we maintained two key orthogonal nucleotides (GU), which replace the UN of the NNyUNAy mammalian branchpoint consensus sequence. The relative strength of the different branchpoints were ranked with SVM_BPfinder (16) in the context of the canonical branchpoint consensus sequence. The sequences were also tested in the orthogonal context to ensure novel cryptic endogenous branchpoint sequences were not created. BP-NC1, which makes two Watson-Crick (WC), three GU and one mismatched base pair with the orthogonal U2 BPRS, had the lowest SVM score while still being identified as a branchpoint.

**Figure 2.**
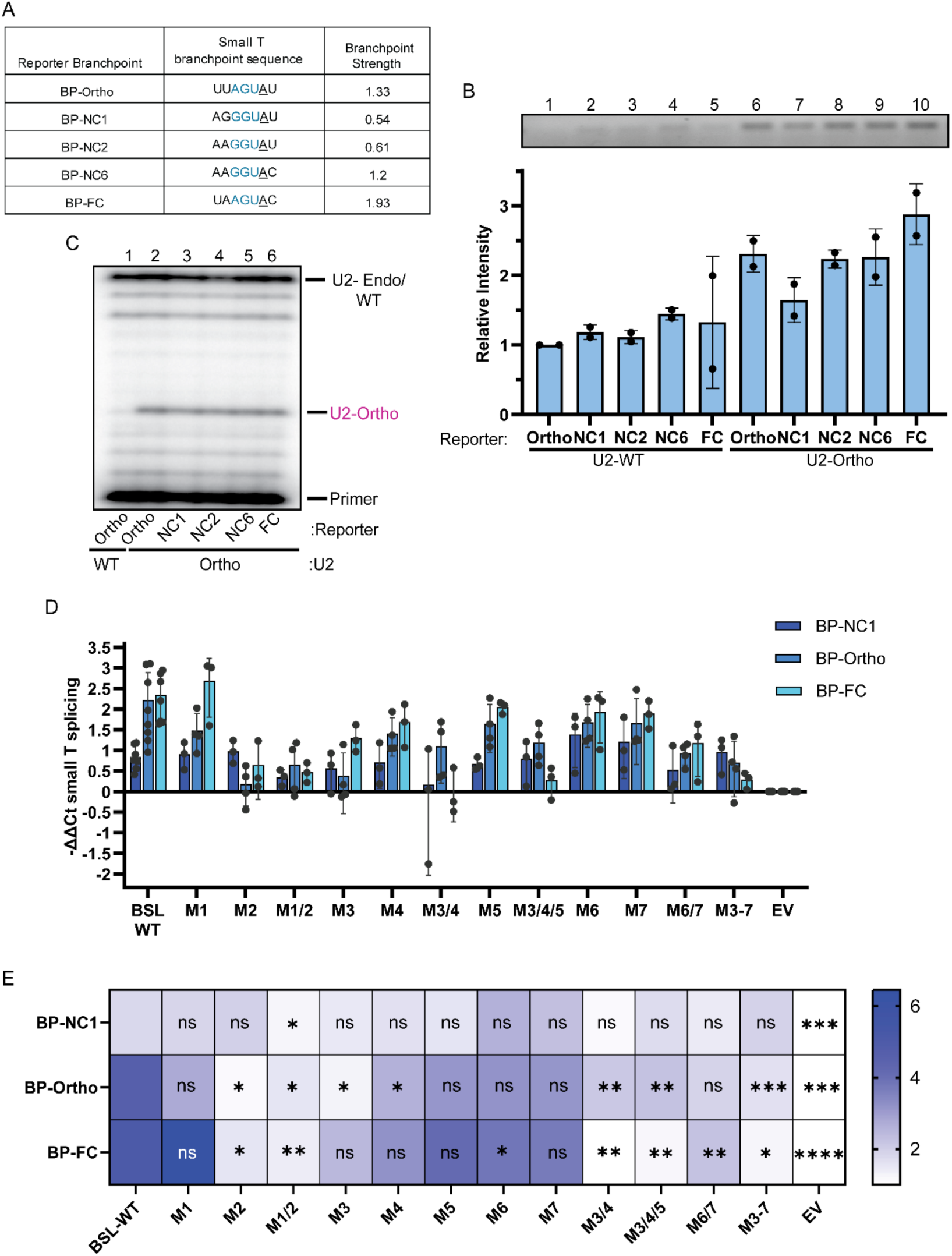
BSL base-pairing potential influences branchpoint sequence recognition. (**A**) Table of orthogonal splicing reporter branchpoints with orthogonal nucleotides in blue. The branchpoint adenosine is underlined. Branchpoint strength is from SVM_BPfinder. (**B**) Small T splicing with different branchpoints. Top: Representative agarose gel of RT-PCR for spliced small T (top). Bottom: Band intensity quantified relative to sample 1 (BP-Ortho and U2-WT); n=2. (**C**) Representative PAGE of radiolabeled U2 snRNA primer extension analysis for samples in (B). Extension stops for endogenous/wildtype U2 snRNA and U2-Ortho are labeled. (**D**) Difference in small T intron splicing by RT-qPCR for different reporter branchpoints with BSL mutants. -ΔΔCt is derived from small T splice junction probe normalized to exon2 and relative to empty vector (EV) of the same transfection batch. BP-Ortho data for Figure 1G and I is included for comparison. (**E**) Data from (D) displayed as a heatmap. In all cases, error bars represent standard deviation.

BP-NC2 and BP-NC6 retain a single mismatch, but incrementally increase the WC base-pairing. BP-Ortho, which was used in Figure 1 experiments, makes five WC and one GU base pairs, and ranks slightly lower than BP-FC, which makes all six WC base pairs. Splicing efficiency of the different reporters correlated with the branchpoint SVM score (Figure 2B, lanes 6-10), with BP-NC1 yielding the lowest level of orthogonal small T splicing and BP-FC yielding the highest when co-transfected with U2-Ortho BSL-WT, which showed consistent expression across all experiments (Figure 2C). Reporter splicing was significantly lower when co-transfected with wildtype U2 snRNA (U2-WT), consistent with the inability of wildtype/endogenous U2 snRNA to recognize the modified orthogonal branchpoint sequences (Figure 2B, lanes 1-5). Together the data suggests that complementarity of the branchpoint to the BPRS of the U2 snRNA is an important factor for efficient recruitment of U2 snRNP to an intron and splicing.

### BSL base-pairing potential influences branchpoint sequence recognition

In *S. cerevisiae*, increasing the base-pairing potential of the BSL decreases the stringency of branchpoint sequence recognition (12). To test whether alterations to the human BSL base-pairing also has a relationship to branchpoint sequence recognition, we selected the weakest (BP-NC1) and strongest (BP-FC) reporters to test in the context of our twelve U2-Ortho BSL mutants. Again, U2-Ortho BSL-WT was expressed at around 10% of the total U2 snRNA in the cell and BSL mutant expression trended similarly as previously observed (Supplementary Figure 2A, B, and C). As expected, splicing efficiency of the branchpoint reporters correlates with the complementarity of the branchpoint sequence for U2-Ortho BSL-WT (Figure 2D, E and Supplementary Figure 3B and C).

For BSL-strengthening mutations, BP-FC reporter splicing in the presence of M1 was the same, if not better than, BSL-WT, while the higher complementarity of the FC branchpoint did not rescue splicing for M2. M1/2 again exhibited little to no expression and no reporter splicing (Figure 2D, E and Supplementary Figure 3B). In contrast, several mutations that perturb BSL base-pairing, including M3, M4, M5, M6 and M7, yield BP-FC reporter splicing somewhat lower than BSL-WT, but not at levels sufficient to meet statistical significance, except for M6.

Previously, we noted that the reason for low reporter splicing with M3 was ambiguous, but this result demonstrates that 1) steady-state orthogonal U2 expression as low as 2% of total U2 snRNA is sufficient to confer significant reporter splicing with the strongest branchpoint and 2) that the G25A mutation likely affects both U2 snRNP biogenesis and U2 snRNA’s ability recognize the weaker BP-Ortho branchpoint sequence. With the combined perturbation of BSL structure in M3/4, M3/4/5 and M3-7, the loss of splicing with BP-FC is dramatic, while U2-Ortho expression levels remain above that of M3. Similarly, the small reduction of BP-FC splicing with M6/7 is not explained by low U2-Ortho expression.

Across all the BSL mutants, splicing levels of the weak BP-NC1 reporter was largely equivalent to U2-Ortho BSL-WT, although its already low splicing level likely does not provide significant range to tease out the mutational effects (Figure 2D, E and Supplementary Figure 3C). None of the BSL mutants show a significant increase in splicing levels compared to BSL-WT, except for M6 and M7 which show a slight but non-significant increase in splicing.

Interestingly, mutants M2 and M3-7, which show virtually no splicing with the stronger branchpoint reporters, yield splicing levels of BP-NC1 comparable to BSL-WT. Together this data indicates that opening near the loop of the BSL may provide some benefit to recognition of low complementarity branchpoint sequences.

Altogether, this data indicates several points: 1) that the BSL is necessary for proper U2 snRNP assembly and splicing, 2) that minor stabilization of the BSL is tolerated but stabilization near the base of the stem, even though functional, along with a fully stabilized BSL, likely prevent proper U2 snRNP assembly and splicing by preventing unwinding of the BSL even at fully complementary branchpoints, 3) BSL requires an intact stem base for proper intron recognition as, even though the splicing effect of M3 was rescued by splicing with a fully complementary branchpoint sequence, increasing the perturbation results in a progressive loss of splicing as complementarity of the branchpoint sequence improves, and 4) the structure of the loop of the BSL is likely important for splicing as M6 and M7 shows no improvement in splicing relative to branchpoint sequence complementarity.

### Compensatory mutations do not reverse the negative effect of some BSL destabilizing mutations

Because many U2 snRNA nucleotides participate in more than one interaction during the splicing process, the effect of mutations can be difficult to parse. While BSL structure may mediate the splicing defect observed with U2-Ortho constructs containing M3 and M4, the altered nucleotides interact with U6 snRNA in Helix Ia, which forms upon catalytic activation of the spliceosome (36). Specifically, M3 changes a G-U base pair to A-U and M4 converts a C-G to U-G. Although modeling these changes appears compatible with the U2/U6 Helix Ia structure, we cannot rule out the potential impact later in the splicing cycle (Supplementary Figure 4B).

To ascertain whether the negative effects of these mutations on reporter splicing can be fully attributed to perturbation of BSL base-pairing, we tested compensatory mutations. M3/M8 (G25A, C45U) and M4/M9 (C28U, G42A) reinstates the base pairs disrupted by M3 and M4 respectively, while M3/8/4/9 reinstates both (Figure 3A). We transfected these constructs with the BP-Ortho reporter into cells and assessed U2-Ortho levels by primer extension and reporter splicing by RT-qPCR. For these experiments, we included new replicates of M3, M4 and M3/M4 for comparison. While U2-Ortho levels remained similar for M4 (partially reduced) M3 and M3/M4 (significantly reduced) relative to BSL-WT, little to no expression of the M3/8, M4/9 and M3/8/4/9 compensatory mutants was detected (Figure 3B). Consistent with the very low expression, reporter splicing with the compensatory mutants was comparable to empty vector control, while the single BSL mutations maintained splicing levels akin to our previous results (Figure 3C). These results indicate that the negative effects of M3 and M4 mutations on U2-Ortho expression and reporter splicing is not simply due the loss of BSL base-pairing, but that nucleotide identity at the altered positions also contributes. Because of highly deleterious effect of the compensatory mutations on U2-Ortho levels, we cannot rule out perturbation of U2/U6 Helix Ia as a contributing factor for the splicing defect associated with M3, M4 and M3/4.

**Figure 3.**
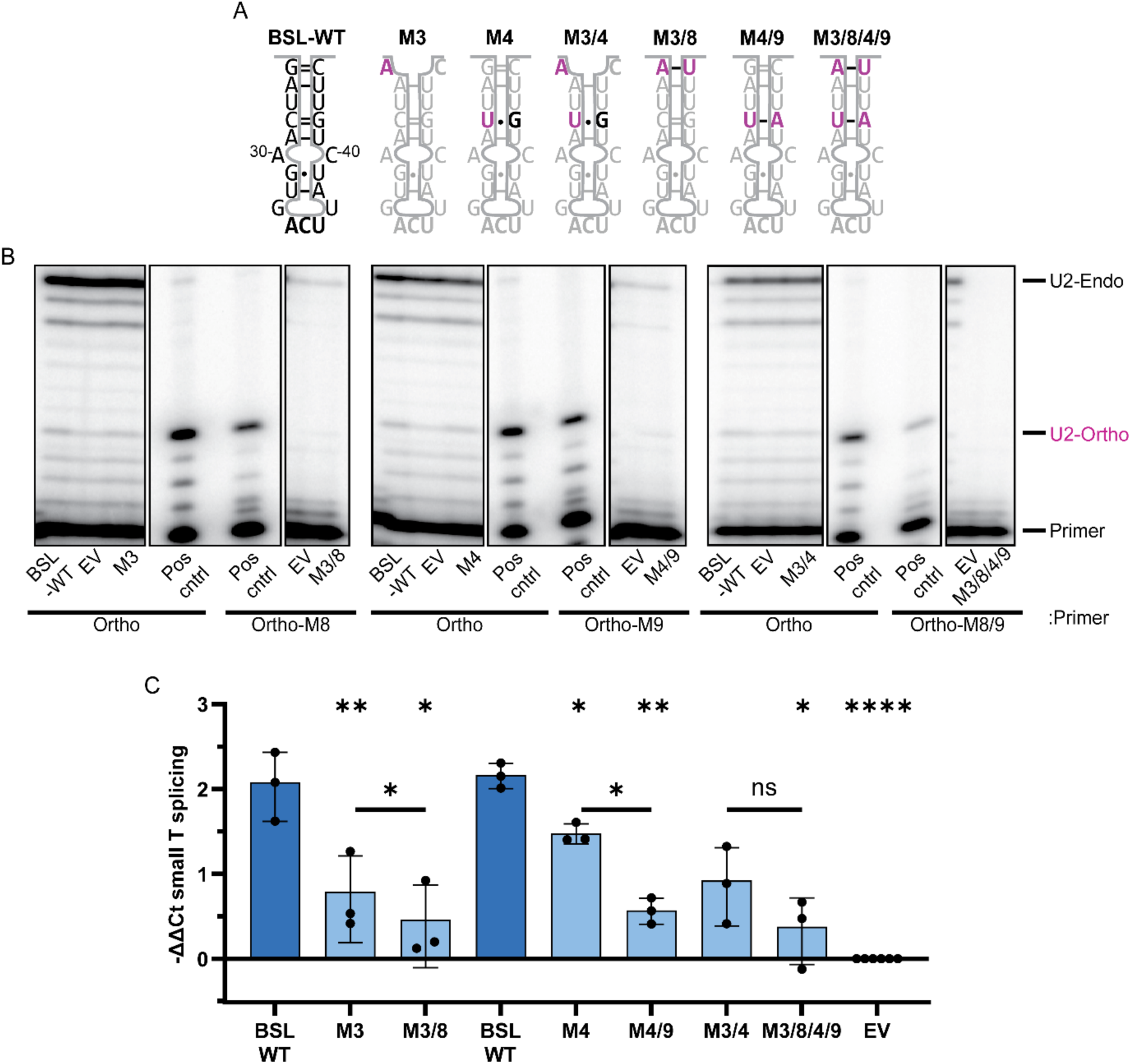
Compensatory mutations do not reverse the negative effect of some BSL weakening mutations. (**A**) Schematic of orthogonal human U2 snRNA BSL wildtype and mutant sequences with predicted base-pairing. Mutations are indicated in pink. (**B**) Representative PAGE of radiolabeled U2 snRNA primer extension analysis for the indicated BSL mutants. Extension stops for endogenous/wildtype U2 snRNA and U2-Ortho are labeled. (**C**) Difference in small T intron splicing by RT-qPCR for samples shown in (B). -ΔΔCt is derived from small T splice junction probe normalized to exon2 and relative to empty vector (EV). Error bars represent standard deviation.

### Expression of U2 snRNA BSL mutations results in global alterations to splicing and gene expression

Recently, mutations to the U2 snRNA RNU2-1 and RNU2-2 BSL region have been associated with several tumor types and neurodevelopmental disorders, including our M4 mutation C28U (13, 37–40). To determine the impact of M4 on other splicing events, we measured the effect on splicing and gene expression in human cells. We were also interested in determining whether the presence of U2 snRNA with a dysfunctional BPRS (*i.e.* U2-Ortho) would have a unique effect on cellular splicing. For these studies, we infected HEK293T cells with lentivirus containing the wildtype RNU2-1 expression locus harboring C28T (U2-C28U), TAG36ACT (U2-UCA) along with an empty lentivirus vector control and selected for stable integration. We extracted total RNA from the cell lines and, after verifying expression of the mutant U2 snRNA by primer extension (Supplementary Figure 5A and B), submitted samples for poly-A selected RNA sequencing. The resulting mapped reads were analyzed for gene expression and splicing changes with the U2 snRNA mutants relative to empty vector control.

Our analysis did not show evidence of widespread splicing changes with either mutant. Only 175 splicing events were significantly different from the empty vector control (FDR ΔPSI ≤ 0.05) in the U2-C28U mutant and 186 events in the U2-UCA mutant (Figure 4A, and Supplementary Tables 2 and 3). While skipped exon (SE) was the most common splicing change, other categories including alternative 3’ splice site (A3), alternative 5’ splice site (A5), alternative first exon (AF), alternative last exon (AL), multiple skipped exons (MS), mutually exclusive exons (MX), and retained introns (RI) were identified. Notably, ∼25% of altered events were shared between the different U2 mutants, suggesting that their impact on splicing is similar (Figure 4B). In several cases, the splicing change involved alternative inclusion of exons containing a premature termination codon (PTC) that targets the transcript for degradation by nonsense mediated decay (NMD). For example, we observed higher skipping of a ‘poison’ cassette exon in splicing factor SRSF6 transcripts and the IDX cassette exon of the HRAS proto-oncogene (Figure 4C and D). Both U2 mutant cell lines also show decreased usage of the alternative 3’ splice site with a downstream PTC in the snRNA 3’-tail processing gene TOE1 transcripts (Figure 4E). Notably, the expected correlated increase in expression of these genes, and thus higher expression of the functional isoform, was confirmed by the DESeq2 analysis described below (Figure 4F). Non-NMD related alternative splicing changes were also identified including a novel functional isoform of the ubiquitin-activating enzyme gene UBA2 with the in-frame skipping of exon 13 (41), and a multiple skipping event of exons 7-9 in the tryptophan metabolizing enzyme gene AFMID that is upregulated in several cancer lines (42) (Figure 4G and H).

**Figure 4:**
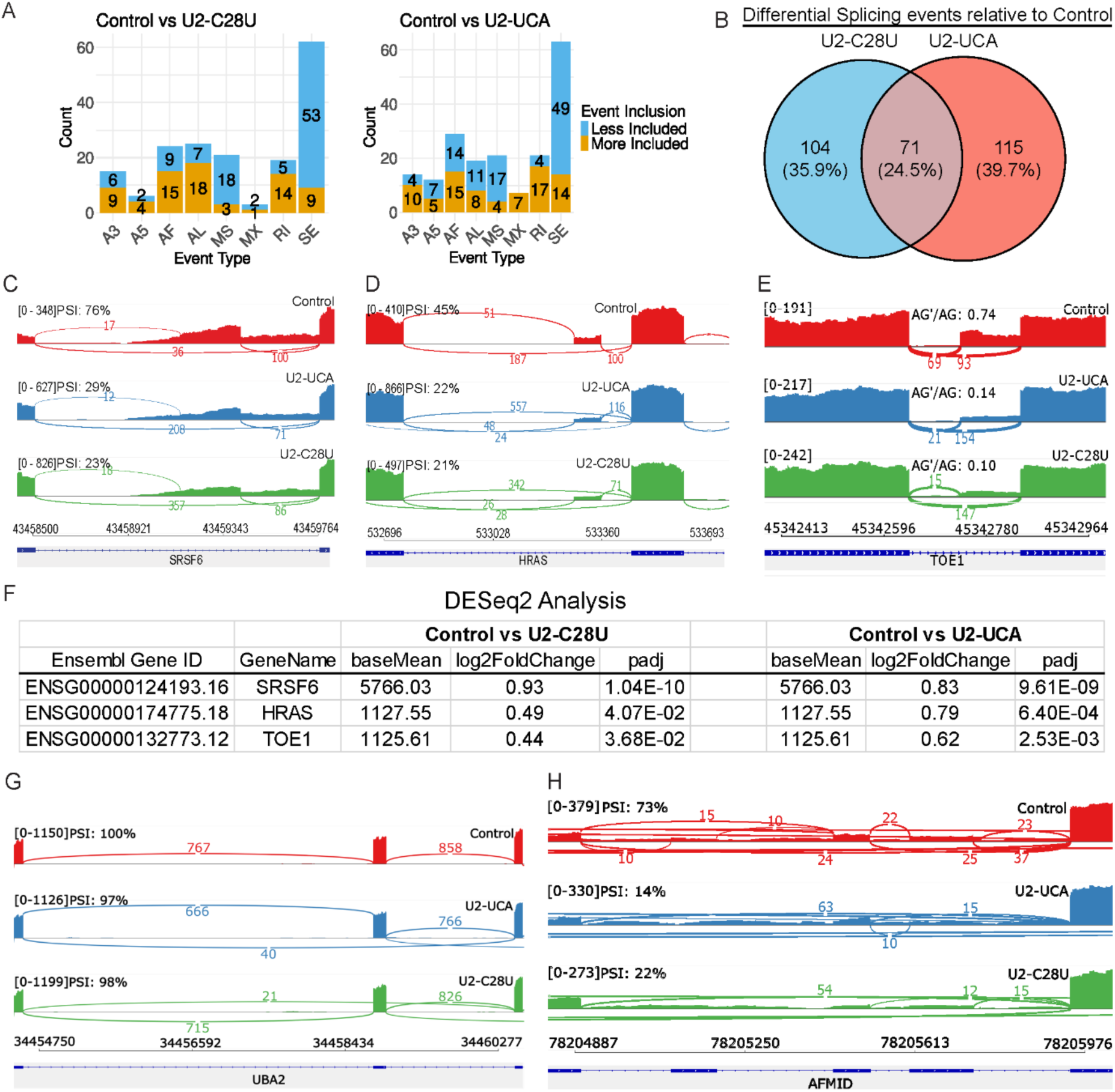
Global splicing changes in cells expressing U2 snRNA BSL or BPRS mutations. (**A**) Bar graph quantifying different types of differential alternative splicing events for each U2 snRNA mutant relative to empty vector control cell line. A3 = Alternative 3’ splice site; A5 = Alternative 5’ splice site; AF = Alternative First Exon; AL = Alternative Last Exon; MS = Multiple Skipped Exons; MX = Mutually Exclusive Exons; RI = Retained Intron; SE = Skipped Exon. (**B**) Venn diagram illustrating differential alternative splicing events overlapping between C28U and U2-UCA. (**C**) Sashimi plot of SE event around exon 3 of SRSF6. Coverage is shown as raw read counts from BAM files and PSI is defined as event inclusion counts/total event counts. (**D**) Sashimi plot of SE event around IDX exon of HRAS. (**E**) Sashimi plot of A3 event at exon 7 of TOE1. (**F**) Differential expression values for SRSF6, HRAS, and TOE1 from DESeq2. (**G**) Sashimi plot of SE event around exon 13 of UBA2. (**H**) Sashimi plot of MS event around exons 7-9 of AFMID.

Much broader effects of both U2 snRNA variants were revealed by differential gene expression analysis. Relative to the empty vector control, there were 1823 down-regulated and 2669 upregulated genes in the U2-C28U cells, and 1923 down-regulated and 2075 upregulated genes in the U2-UCA cells (Figure 5A and B). Like the splicing changes, many differentially expressed genes (∼47%) were shared between the two cell lines (Figure 5C). However, only ∼30% of genes with altered splicing also showed differential expression with both U2 snRNA variants (Figure 5D and E).

**Figure 5:**
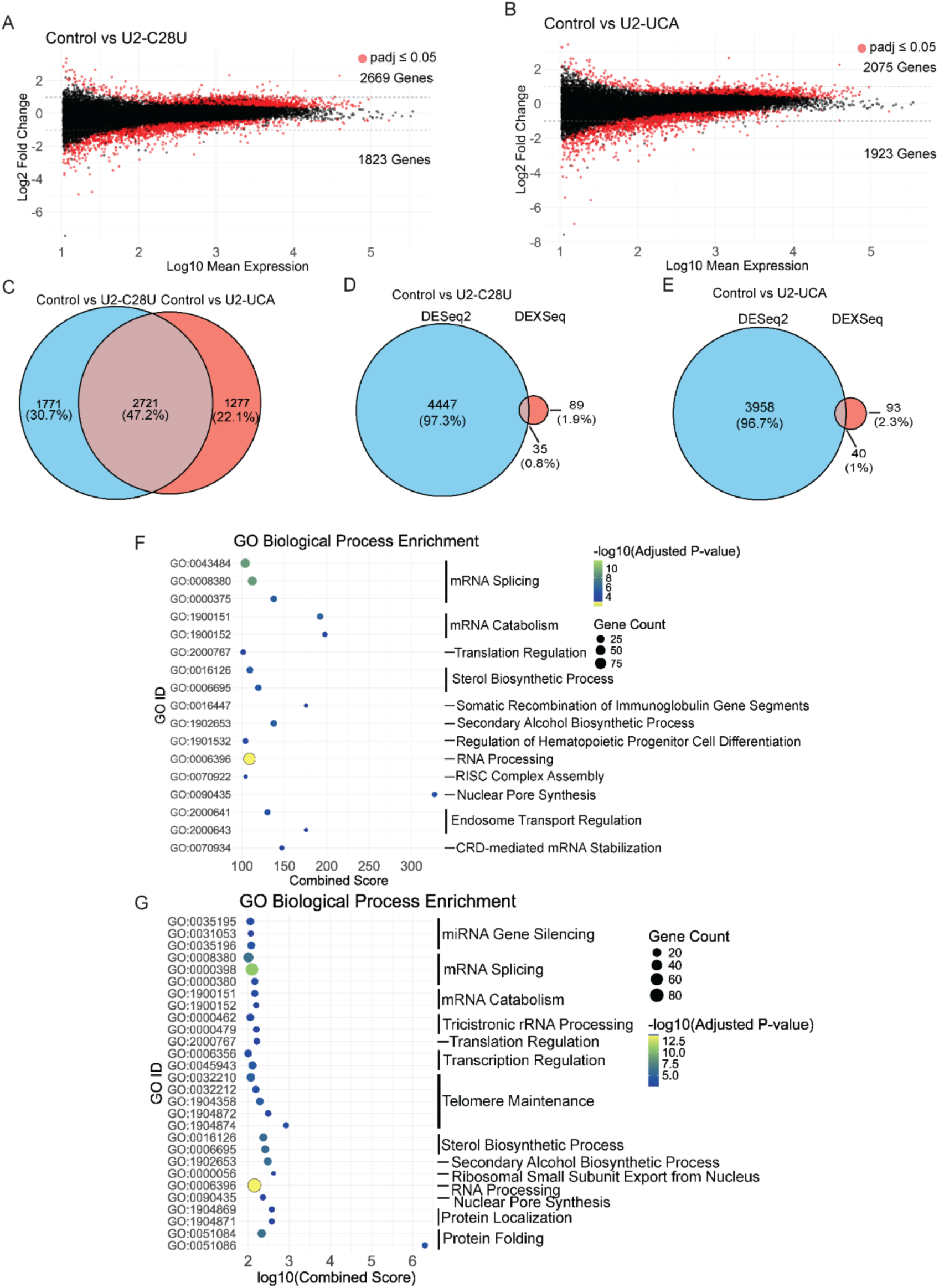
U2 snRNA BSL and BPRS mutations results in global alterations to gene expression. (**A**) Plot of log2 Fold Change versus Log10 Mean Expression for genes from Control vs U2-C28U comparison. Number of significant upregulated and downregulated genes (padj ≤ 0.05) is shown. (**B**) Same as (A) for Control vs U2-UCA comparison. (**C**) Venn diagram illustrating differentially expressed genes overlapping between U2-C28U and U2-UCA. (**D**) Venn diagram illustrating overlap of genes with differentially expression and alternative splicing for U2-C28U. (**E**) Same as (D) for U2-UCA. (**F**) Bubble plot of GO Biological Process terms enriched in significantly upregulated genes from U2-C28U grouped by similar pathways. Color represents the significance of the enrichment (-log10(adjusted *p*-value)) and bubble size indicates the number of genes identified. (**G**) Same as (F) for U2-UCA.

Gene ontology (GO) Biological Process enrichment analysis of both the differentially spliced and the downregulated genes identified no significantly enriched terms (adjusted *p*-value ≤ 0.05) in either cell line. In contrast, several pathways showed enrichment in the upregulated genes in both cell lines, including many pathways linked to general regulation of gene expression for RNA processing, translation, RNA decay, and nuclear pore formation (Figure 5F and G). The magnitude of the expression changes for many of these genes was relatively modest. For example, the upregulation of genes involved in RNA processing is between 0.3-1 Log_2_ fold change. However, these genes are normally well-expressed (>2.5 Log_10_ fold mean expression), and cells may have limited ability to upregulate them much further (Supplementary Tables 4 and 5).

Overall, this analysis indicates that the presence of an altered U2 snRNA can be tolerated in cells with minimal impact on RNA splicing choices. However, there is an effect on gene expression. We postulate that the mutant U2 snRNPs may compete with WT U2 snRNP to slightly decrease splicing efficiency broadly. For many genes, this increases RNA decay of the transcripts through pathways like NMD. Cells may also have mechanisms to respond to the decrease in splicing efficiency by upregulating genes involved in RNA processing and gene expression more generally.

How mutant U2 snRNA identified in cancers contributes to the disease is unknown. Considering that the overexpression of the oncogene MYC is also associated with upregulation of splicing factors (43) and increased skipping of the HRAS IDX cassette exon (44), we looked more closely at its expression. The MYC transcript levels in both U2 mutant cell line compared to empty vector control were not significantly different. However, GO term analysis of upregulated genes in the U2 mutants using the Molecular Signatures Database (MSigDB) Hallmark sets revealed that MYC target genes, mTORC signaling genes and G2-M checkpoint genes are highly overrepresented (Figure 6). The MYC targets, like the RNA splicing and RNA processing genes, showed modest expression change (between 0.3 and 1 Log_2_ fold change) supported high read counts (>1.7 Log10 Mean Expression) (Supplementary Tables 4 and 5). Importantly, most genes identified as MYC targets were not associated with splicing or the spliceosome, indicating that the effect of mutant U2 snRNA expression extends past dysregulation of pre-mRNA processing pathways and may contribute to cancer development or progression by phenocopying MYC overexpression.

**Figure 6.**
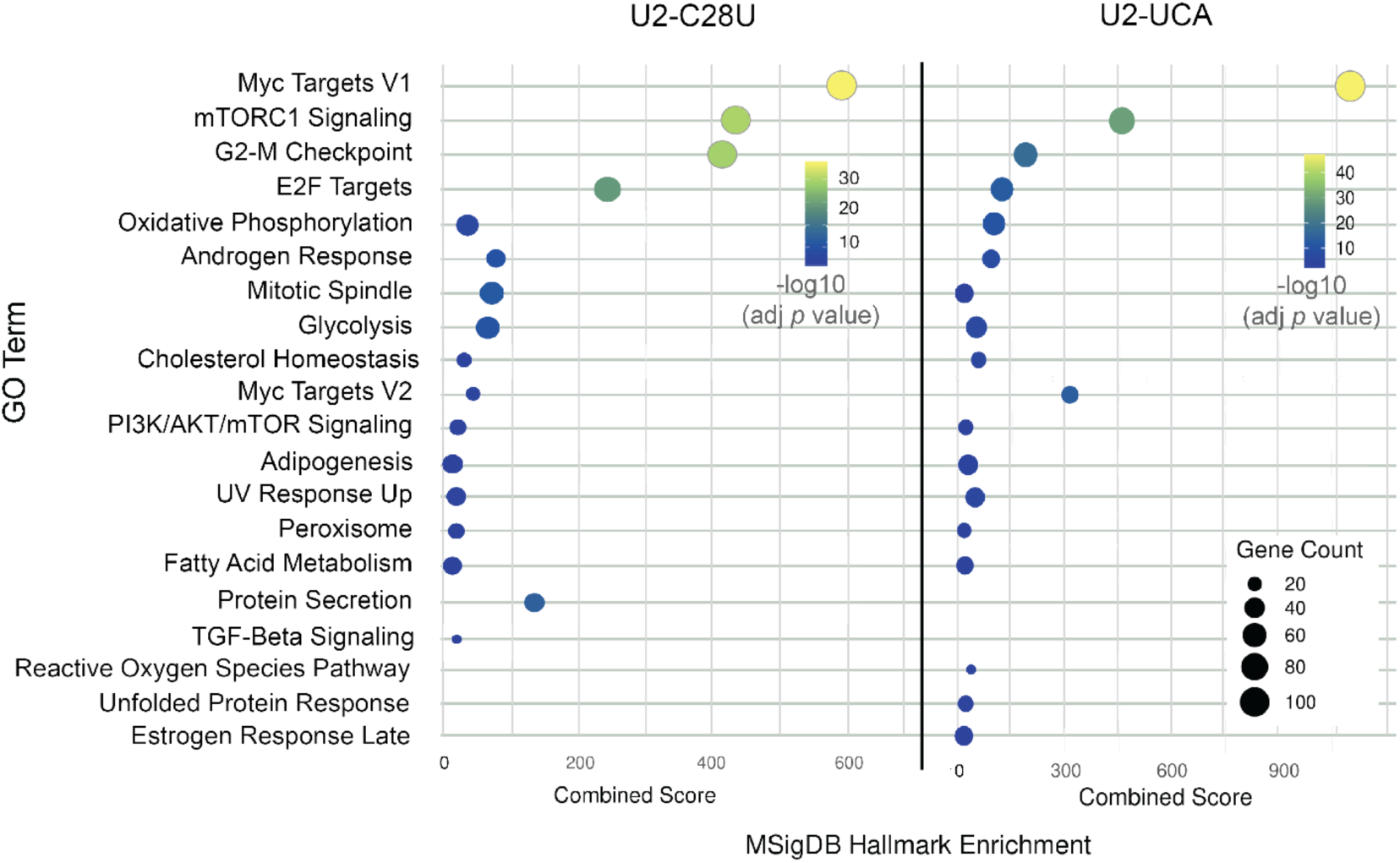
MYC-target genes are upregulated with mutant U2 snRNA expression. Bubble plot of MSigDB Hallmark Enrichment GO terms enriched for significantly upregulated genes with U2-C28U and U2-UCA. Color represents the significance of the enrichment (-log10(adjusted *p*-value)) and bubble size indicates the number of genes identified.

## DISCUSSION

U2 snRNA is a highly dynamic molecule with several competing structures, both internally (*e.g.* Stem IIa vs. Stem IIc) and externally (*e.g.* Stem I vs. U2/U6 Stem II). Based on studies in yeast, the BSL was first described as a conserved and functional feature of the U2 snRNA (12), and its structure was later confirmed in the cryo-EM model of the human U2 snRNP, although it was not well resolved (9–11). In the absence of proteins, U2 snRNA does not adopt the BSL structure (45), meaning that BSL formation is controlled by the U2 snRNP proteins that surround it. Furthermore, the BSL is mutually exclusive with the branch helix, indicating that BSL unwinding is a regulatory step of spliceosome assembly. Altered base-pairing within the BSL would therefore be predicted to influence both U2 snRNP’s assembly, structure and branchpoint sequence recognition. Because of the many constraints against manipulating U2 snRNA genomic sequence, we used an orthogonal splicing system to carry out a mutational analysis of the human U2 snRNA BSL sequence. In the following paragraphs we discuss how the results support a role for the BSL in regulating branchpoint sequence recognition and branch helix formation, and how proper BSL structure is vital for stable U2 snRNA expression (Figure 7A and B).

**Figure 7.**
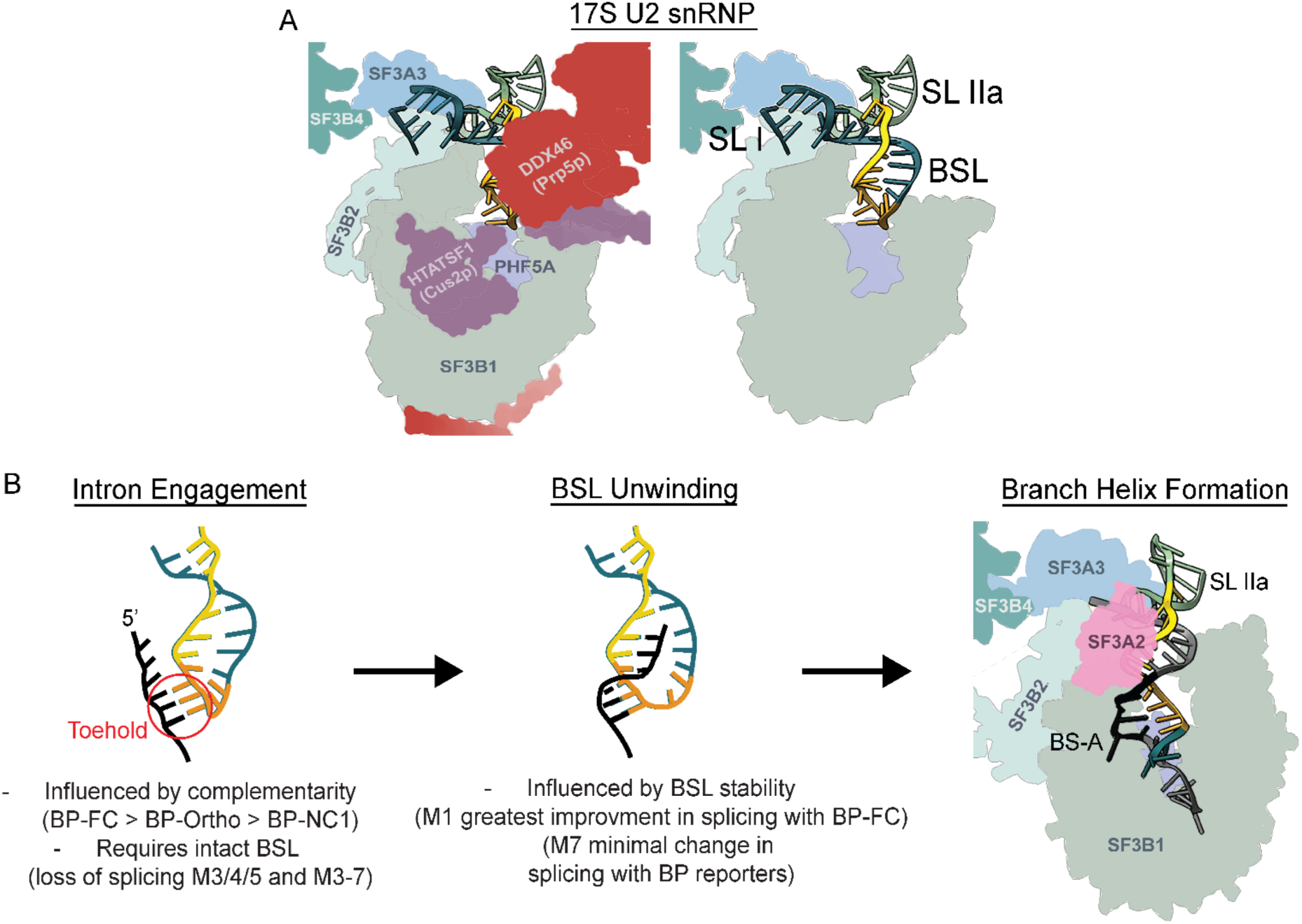
Branchpoint sequence complementarity and BSL base-pairing regulate branch helix formation. (**A**) Schematic of BSL and surrounding proteins before intron engagement based on cryo-EM models of 17S U2 snRNP (PDB 6Y50, 7EVO). (Left) Proteins in the vicinity of the BSL are labeled. (Right) Proteins HTATSF1 and DDX46 are removed to better view the structure of SLI, BSL and SLIIa of the U2 snRNA. Nucleotides that contact the intron in BPRS and BSL are colored orange and yellow respectively. (**B**) Model for BSL regulation of branch helix formation. (Intron Engagement) The intron region containing the branchpoint sequence is brought close to the BSL and the toehold is formed by base-pairing between nucleotides 35-AGΨ of the U2 snRNA and NyU of the intron branchpoint sequence. (BSL Unwinding) Invasion of the intron and formation of additional base pairs between the intron and BSL induces BSL unwinding. (Branch Helix Formation) Complete unwinding of the BSL allows for formation of the rest of the branch helix. The bulged branchpoint adenosine is captured by SF3B1.

Prior to branch helix formation, the U2 snRNA is brought to the vicinity of the branchpoint sequence with the BSL intact and with nucleotides 35-37 in the BSL loop available for initial recognition by the branchpoint sequence (Figure 7B). These nucleotides base pair with the NyU of the human consensus, NNyUNAy (7, 8). The impact of BSL mutations on splicing of orthogonal reporters harboring different branchpoint sequences indicates that the complementarity regulates initial branchpoint sequence recognition and induces BSL unwinding (Figure 7B). This finding expands a model of branch helix formation proposed by Pena and coworkers based on their structure of an SF3B1 inhibitor-stalled spliceosome with a partially formed branch helix and full extended helix (46). They postulated that intron base-pairing with the BSL loop establishes a ”toehold” and then further intron/U2 base-pairing displaces BSL interactions to form the extended branch helix. This model predicts that the strength of the toehold interaction would therefore dictate the efficiency of strand invasion by the intron (47–49). Consistent with this idea, we see a difference in splicing efficiency when U2 snRNA complementarity within the toehold is altered. In the orthogonal system, U2-Ortho CUU is fully complementary to BP-FC, which has the highest splicing efficiency, while the single WC base pair and two G-U base pairs with NC-1 results in much lower splicing (Figure 7B).

Because intron base-pairing in the extended helix should contribute to BSL unwinding, the model also predicts that a weaker BSL near the toehold would be less dependent on the strength of the toehold to initiate unwinding, while increased BSL base-pairing would be more dependent on perfect base-pairing. Consistent with these predictions, when there are fewer BSL WC base pairs adjacent to the toehold (M6 and M7), splicing efficiency no longer correlates with toehold complementarity (Figure 7B). In contrast, when a WC base pair is added to the BSL (M1) the improvement in splicing efficiency conferred by the fully complementary branchpoint of BP-FC is larger compared to BSL-WT. We propose that the strength of base-pairing within the BSL could serve as a check on toehold complementarity to help maintain branchpoint fidelity. With weaker human branchpoint sequences, we imagine that other factors help extend the lifetime of the toehold to support BSL unwinding.

Several proteins have been linked to BSL structure, fidelity of branchpoint recognition, and branch helix formation. The cryo-EM model of 17S U2 snRNP shows the BSL structure pinned down by U2 snRNA interactions with SF3B1, SF3A3, SF3B2 and HTATSF1 (Figure 7A) (9–11, 46). HTATSF1 (Cus2p in *S. cerevisiae*), cradles the loop of the BSL in the 17S U2 snRNP and limits access to the nucleotides that would form the toehold with an intron. It likely serves as a gatekeeper of BSL until it is released through the ATPase activity of DDX46 (Prp5p in *S. cerevisiae*). Tholen *et al* reported that removal of HTATSF1 in the absence of a competing intron results in a novel and non-productive structure termed the Branch Helix-Mimicking Stem-Loop where nucleotides in the 5’ end of U2 snRNA are base paired with the BPRS and with three BSL nucleotides (11). Our results further underscore the need to determine the relative timing of establishing the toehold, HTATSF1 release, and freeing of the BSL for unwinding to allow for branch helix formation.

In addition to its role in splicing, proper formation of the BSL structure is likely necessary for complete assembly of the U2 snRNP and may present as a biogenesis quality control checkpoint. All U2-Ortho constructs in which the first base pair of the BSL stem is disrupted (M3) consistently showed decreased stable expression relative to BSL-WT, which may reflect a requirement for BSL formation during U2 snRNP biogenesis. Notably, a tryptophan side chain from SF3A3 stacks with G25 in the G25-C45 base pair (9–11). This interaction could be necessary for nucleating BSL formation, positioning the BSL within the 17S U2 snRNP, or docking of SF3A3 into the U2 snRNP, any of which could result in the dramatic phenotypes we observed. Increased WC base-pairing in the middle of the stem is also not tolerated (M2, and M1/2) indicating that a competing structure may also be important for forming the U2 snRNP. However, another possibility is that the impact on stability is related to RNA modification because nucleotide U43 is normally pseudouridylated. U2 snRNA is highly modified, including within the BSL (50–52). Pseudouridylation was shown to important for full U2 snRNP assembly and splicing function in frog oocytes (51). It is possible that some U2 snRNA modifications are monitored to ensure proper snRNP assembly, so that improperly folded or unmodified U2 snRNA is degraded. While quality control mechanisms for 3’ end formation and SM protein association exist for snRNPs(53, 54) to our knowledge, no quality-control mechanism for other U2 snRNA structures in splicing has been reported. Our data motivate further investigation of these questions.

Bousquets-Muñoz and collegues identified specific U2 snRNA mutations in several cancers including mantle cell lymphoma, chronic lymphocytic leukemia, B-cell non-Hodgkin lymphoma, prostate adenocarcinoma, and pancreatic cancer, although in comparison to matched normal samples, no significantly differentially spliced or expressed genes were observed (13). We find that expression of the C28U U2 snRNA mutant in HEK293T cells results in modest but significant splicing changes and a wider impact on gene expression. How these changes could contribute to carcinogenesis is not known, but it is tempting to speculate they would be similar to the changes associated with well-documented cancer mutations in other U2-snRNP proteins. We do not, however, observe the verified differences in alternative 3’ splice site selection identified with SF3B1 mutations in patient samples and engineered cell lines (55–61). Moreover, genes reported as downregulated with mutant SF3B1 are upregulated with C28U U2 snRNA mutation. We also do not see reported splicing changes associated with U2AF1 and SRSF2 cancer mutations (62–68). On the other hand, a recent study examining the effect of both C28U and C28A mutations in RNU2-1 and RNU2-2 sequences expressed from the RNU2-2 single locus reported findings similar to ours (69). Transcriptomic analysis of a hotspot mutation in the U1 snRNA 5’ splice site recognition sequence revealed upregulation of genes in the same pre-mRNA processing pathways affected by the C28U U2 snRNA cancer mutation (70, 71). Interestingly, the expression of a U2 snRNA with a mutant BRPS, which is not expected to be functional for normal splicing, results in very similar changes. We favor the possibility that the mild splicing defects produced by the presence of a relatively small population of mutant snRNAs results in a cellular response that generally upregulates the machinery involved in gene expression and cell proliferation. Further studies to separate direct and indirect effects of mutant U2 snRNA expression may illuminate strategies to target the mutations as a potential cancer therapy.

## DATA AVAILABILITY

The underlying data of this study are available in the article and supplementary material. The RNA-sequencing data are available in Gene Expression Omnibus (GEO) database at https://www.ncbi.nlm.nih.gov/geo/, under accession number GSE303759. UCSC genome browser tracks of the DESeq2 size factor normalized read counts can be found at: https://genome.ucsc.edu/s/mstevers/U2muttracks_groupedcolor.

Supplementary Data are available at NAR Online.

## AUTHOR CONTRIBUTIONS

M.B.S. Conceptualization, Data Curation, Formal Analysis, Funding Acquisition, Investigation, Methodology, Project Administration, Validation, Visualization, Writing – original draft, Writing – review and editing. S.K. Investigation, Software, Resources. M.S.J. Conceptualization, Funding Acquisition, Methodology, Project Administration, Supervision, Visualization, Writing – review and editing.

## ACKNOWLEDGEMENTS

We thank Dr. Manuel Ares and Dr. Hannah Maul-Newby for insightful discussion and comments on the manuscript. We thank our many colleagues at UCSC for assistance, especially Dr. Victor Tse and Dr. Zach Neeb (RT-qPCR assays) and Dr. Angela Brooks and Dr. Cindy Liang (RNA sequencing) and Kristina Oh (cloning U2 snRNA mutant plasmids). RNA sequencing was carried out at the DNA Technologies and Expression Analysis Cores at the UC Davis Genome Center, supported by NIH Shared Instrumentation Grant 1S10OD010786-01. Lastly, we thank Dr. Nicholas Stevers at the University of California San Francisco for lentivirus reagents and protocol, computational resources and RNA-seq analysis support.

## REFERENCES

1. Reed, R. and Maniatis, T. (1985) Intron sequences involved in lariat formation during pre-mRNA splicing. Cell, 41, 95–105.

2. Parker, R., Siliciano, P.G. and Guthrie, C. (1987) Recognition of the TACTAAC box during mRNA splicing in yeast involves base pairing to the U2-like snRNA. Cell, 49, 229–239.

3. Zhuang, Y. and Weiner, A.M. (1989) A compensatory base change in human U2 snRNA can suppress a branch site mutation. Genes Dev., 3, 1545–1552.

4. Wu, J. and Manley, J.L. (1989) Mammalian pre-mRNA branch site selection by U2 snRNP involves base pairing. Genes Dev., 3, 1553–1561.

5. Li, M. and Pritchard, P.H. (2000) Characterization of the Effects of Mutations in the Putative Branchpoint Sequence of Intron 4 on the Splicing within the Human Lecithin:cholesterol Acyltransferase Gene. Journal of Biological Chemistry, 275, 18079–18084.

6. Královičová, J., Houngninou-Molango, S., Krämer, A. and Vořechovský, I. (2004) Branch site haplotypes that control alternative splicing. Human Molecular Genetics, 13, 3189–3202.

7. Gao, K., Masuda, A., Matsuura, T. and Ohno, K. (2008) Human branch point consensus sequence is yUnAy. Nucleic Acids Research, 36, 2257–2267.

8. Mercer, T.R., Clark, M.B., Andersen, S.B., Brunck, M.E., Haerty, W., Crawford, J., Taft, R.J., Nielsen, L.K., Dinger, M.E. and Mattick, J.S. (2015) Genome-wide discovery of human splicing branchpoints. Genome Res., 25, 290–303.

9. Zhang, Z., Will, C.L., Bertram, K., Dybkov, O., Hartmuth, K., Agafonov, D.E., Hofele, R., Urlaub, H., Kastner, B., Lührmann, R., et al. (2020) Molecular architecture of the human 17S U2 snRNP. Nature, 583, 310–313.

10. Zhang, X., Zhan, X., Bian, T., Yang, F., Li, P., Lu, Y., Xing, Z., Fan, R., Zhang, Q.C. and Shi, Y. (2024) Structural insights into branch site proofreading by human spliceosome. Nat Struct Mol Biol, 31, 835–845.

11. Tholen, J., Razew, M., Weis, F. and Galej, W.P. (2021) Structural basis of branch site recognition by the human spliceosome.

12. Perriman, R. and Ares, M. (2010) Invariant U2 snRNA Nucleotides Form a Stem Loop to Recognize the Intron Early in Splicing. Molecular Cell, 38, 416–427.

13. Bousquets-Muñoz, P., Díaz-Navarro, A., Nadeu, F., Sánchez-Pitiot, A., López-Tamargo, S., Shuai, S., Balbín, M., Tubio, J.M.C., Beà, S., Martin-Subero, J.I., et al. (2022) PanCancer analysis of somatic mutations in repetitive regions reveals recurrent mutations in snRNA U2. *npj Genom*. Med., 7, 19.

14. WUt, J. and Manley, J.L. (1992) Multiple Functional Domains of Human U2 Small Nuclear RNA: Strengthening Conserved Stem I Can Block Splicing. 12, 10.

15. Ares, M., Mangin, M. and Weiner, A.M. (1985) Orientation-Dependent Transcriptional Activator Upstream of a Human U2 snRNA Gene. MOL. CELL. BIOL., 5.

16. Corvelo, A., Hallegger, M., Smith, C.W.J. and Eyras, E. (2010) Genome-Wide Association between Branch Point Properties and Alternative Splicing. PLoS Comput Biol, 6, e1001016.

17. Camacho Londoño, J. and Philipp, S.E. (2016) A reliable method for quantification of splice variants using RT-qPCR. BMC Molecular Biol, 17, 8.

18. Patro, R., Duggal, G., Love, M.I., Irizarry, R.A. and Kingsford, C. (2017) Salmon provides fast and bias-aware quantification of transcript expression. Nat Methods, 14, 417–419.

19. Ewels, P., Magnusson, M., Lundin, S. and Käller, M. (2016) MultiQC: summarize analysis results for multiple tools and samples in a single report. Bioinformatics, 32, 3047–3048.

20. Andrews, S. (2010) FastQC: a quality control tool for high throughput sequence data.

21. Krueger, K. Trim galore. Wrapper Tool Cutadapt FastQC Consistently Apply Qual Adapt Trimming FastQ Files.

22. Dobin, A., Davis, C.A., Schlesinger, F., Drenkow, J., Zaleski, C., Jha, S., Batut, P., Chaisson, M. and Gingeras, T.R. (2013) STAR: ultrafast universal RNA-seq aligner. Bioinformatics, 29, 15–21.

23. Li, H., Handsaker, B., Wysoker, A., Fennell, T., Ruan, J., Homer, N., Marth, G., Abecasis, G., Durbin, R., and 1000 Genome Project Data Processing Subgroup (2009) The Sequence Alignment/Map format and SAMtools. Bioinformatics, 25, 2078–2079.

24. Ramírez, F., Dündar, F., Diehl, S., Grüning, B.A. and Manke, T. (2014) deepTools: a flexible platform for exploring deep-sequencing data. Nucleic Acids Research, 42, W187–W191.

25. Love, M.I., Huber, W. and Anders, S. (2014) Moderated estimation of fold change and dispersion for RNA-seq data with DESeq2. Genome Biol, 15, 550.

26. Chen, E.Y., Tan, C.M., Kou, Y., Duan, Q., Wang, Z., Meirelles, G.V., Clark, N.R. and Ma’ayan, A. (2013) Enrichr: interactive and collaborative HTML5 gene list enrichment analysis tool. BMC Bioinformatics, 14, 128.

27. Kuleshov, M.V., Jones, M.R., Rouillard, A.D., Fernandez, N.F., Duan, Q., Wang, Z., Koplev, S., Jenkins, S.L., Jagodnik, K.M., Lachmann, A., et al. (2016) Enrichr: a comprehensive gene set enrichment analysis web server 2016 update. Nucleic Acids Res, 44, W90–W97.

28. Xie, Z., Bailey, A., Kuleshov, M.V., Clarke, D.J.B., Evangelista, J.E., Jenkins, S.L., Lachmann, A., Wojciechowicz, M.L., Kropiwnicki, E., Jagodnik, K.M., et al. (2021) Gene Set Knowledge Discovery with Enrichr. Current Protocols, 1, e90.

29. Pertea, M., Pertea, G.M., Antonescu, C.M., Chang, T.-C., Mendell, J.T. and Salzberg, S.L. (2015) StringTie enables improved reconstruction of a transcriptome from RNA-seq reads. Nat Biotechnol, 33, 290–295.

30. Ritter, A.J., Wallace, A., Ronaghi, N. and Sanford, J.R. (2024) junctionCounts: comprehensive alternative splicing analysis and prediction of isoform-level impacts to the coding sequence. NAR Genomics and Bioinformatics, 6, lqae093.

31. Robinson, J.T. (2011) Integrative genomics viewer. correspondence, 29.

32. Pavelitz, T., Rusché, L., Matera, A.G., Scharf, J.M. and Weiner, A.M. (1995) Concerted evolution of the tandem array encoding primate U2 snRNA occurs in situ, without changing the cytological context of the RNU2 locus. The EMBO Journal, 14, 169–177.

33. Tessereau, C., Buisson, M., Monnet, N., Imbert, M., Barjhoux, L., Schluth-Bolard, C., Sanlaville, D., Conseiller, E., Ceppi, M., Sinilnikova, O.M., et al. (2013) Direct Visualization of the Highly Polymorphic RNU2 Locus in Proximity to the BRCA1 Gene. PLoS ONE, 8, e76054.

34. Arsdell, S.W.V. and Weiner, A.M. (1984) Human genes for U2 small nuclear RNA are tandemly repeated. 4, 8.

35. Mabin, J.W., Lewis, P.W., Brow, D.A. and Dvinge, H. (2021) Human spliceosomal snRNA sequence variants generate variant spliceosomes. RNA, 27, 1186–1203.

36. Haselbach, D., Komarov, I., Agafonov, D.E., Hartmuth, K., Graf, B., Dybkov, O., Urlaub, H., Kastner, B., Lührmann, R. and Stark, H. (2018) Structure and Conformational Dynamics of the Human Spliceosomal Bact Complex. Cell, 172, 454–464.e11.

37. Greene, D., De Wispelaere, K., Lees, J., Codina-Solà, M., Jensson, B.O., Hales, E., Katrinecz, A., Nieto Molina, E., Pascoal, S., Pfundt, R., et al. (2025) Mutations in the small nuclear RNA gene RNU2-2 cause a severe neurodevelopmental disorder with prominent epilepsy. Nat Genet, 57, 1367–1373.

38. Jia, Y., Mu, J.C. and Ackerman, S.L. (2012) Mutation of a U2 snRNA Gene Causes Global Disruption of Alternative Splicing and Neurodegeneration. Cell, 148, 296–308.

39. Leitão, E., Santini, A., Cogne, B., Essid, M., Athanasiadou, M., LaFlamme, W., Marijon, P., Bernard, V., Chatron, N., Barcia, G., et al. Systematic analysis of snRNA genes reveals frequent RNU2-2 variants in dominant and recessive developmental and epileptic encephalopathies.

40. Jackson, A., Blakes, A.J., Wall, E., Clarke, N., Agrawal, S., Blair, E., Brady, A.F., Brittain, H., Drinkall, N., Elmslie, F., et al. 1 Biallelic variants in RNU2-2 cause a remarkably 2 frequent developmental epileptic encephalopathy.

41. Chmielarska, K. Biochemical and cell biological characterisation of Sumo E1 activating enzyme Aos1/Uba2.

42. Lin, K.-T., Ma, W.K., Scharner, J., Liu, Y.-R. and Krainer, A.R. (2018) A human-specific switch of alternatively spliced *AFMID* isoforms contributes to *TP53* mutations and tumor recurrence in hepatocellular carcinoma. Genome Res., 28, 275–284.

43. Koh, C.M., Bezzi, M., Low, D.H.P., Ang, W.X., Teo, S.X., Gay, F.P.H., Al-Haddawi, M., Tan, S.Y., Osato, M., Sabò, A., et al. (2015) MYC regulates the core pre-mRNA splicing machinery as an essential step in lymphomagenesis. Nature, 523, 96–100.

44. Chen, X., Yang, H.T., Zhang, B., Phillips, J.W., Cheng, D., Rigo, F., Witte, O.N., Xing, Y. and Black, D.L. (2023) The RNA-binding proteins hnRNP H and F regulate splicing of a MYC-dependent HRAS exon in prostate cancer cells. Proc. Natl. Acad. Sci. U.S.A., 120, e2220190120.

45. Urabe, V.K., Stevers, M., Ghosh, A.K. and Jurica, M.S. (2021) U2 snRNA structure is influenced by SF3A and SF3B proteins but not by SF3B inhibitors. PLoS ONE, 16, e0258551.

46. Cretu, C., Gee, P., Liu, X., Agrawal, A., Nguyen, T.-V., Ghosh, A.K., Cook, A., Jurica, M., Larsen, N.A. and Pena, V. (2021) Structural basis of intron selection by U2 snRNP in the presence of covalent inhibitors. Nat Commun, 12, 4491.

47. Green, S.J., Lubrich, D. and Turberfield, A.J. (2006) DNA Hairpins: Fuel for Autonomous DNA Devices. Biophysical Journal, 91, 2966–2975.

48. Guo, Y., Wei, B., Xiao, S., Yao, D., Li, H., Xu, H., Song, T., Li, X. and Liang, H. (2017) Recent advances in molecular machines based on toehold-mediated strand displacement reaction. Quant. Biol., 5, 25–41.

49. Šulc, P., Ouldridge, T.E., Romano, F., Doye, J.P.K. and Louis, A.A. (2015) Modelling Toehold-Mediated RNA Strand Displacement. Biophysical Journal, 108, 1238–1247.

50. Dönmez, G., Hartmuth, K. and Lührmann, R. (2004) Modified nucleotides at the 5′ end of human U2 snRNA are required for spliceosomal E-complex formation. RNA, 10, 1925– 1933.

51. Zhao, X. and Yu, Y.-T. (2004) Pseudouridines in and near the branch site recognition region of U2 snRNA are required for snRNP biogenesis and pre-mRNA splicing in *Xenopus* oocytes. RNA, 10, 681–690.

52. Yu, Y.-T. (1998) Modifications of U2 snRNA are required for snRNP assembly and pre-mRNA splicing. The EMBO Journal, 17, 5783–5795.

53. Ma, T., Xiong, E.S., Lardelli, R.M. and Lykke-Andersen, J. (2024) Sm complex assembly and 5′ cap trimethylation promote selective processing of snRNAs by the 3′ exonuclease TOE1. Proc. Natl. Acad. Sci. U.S.A., 121, e2315259121.

54. Lardelli, R.M. and Lykke-Andersen, J. (2020) Competition between maturation and degradation drives human snRNA 3′ end quality control. Genes Dev., 34, 989–1001.

55. Pacholewska, A., Lienhard, M., Brüggemann, M., Hänel, H., Bilalli, L., Königs, A., Heß, F., Becker, K., Köhrer, K., Kaiser, J., et al. (2024) Long-read transcriptome sequencing of CLL and MDS patients uncovers molecular effects of *SF3B1* mutations. Genome Res., 34, 1832–1848.

56. Darman, R.B., Seiler, M., Agrawal, A.A., Lim, K.H., Peng, S., Aird, D., Bailey, S.L., Bhavsar, E.B., Chan, B., Colla, S., et al. (2015) Cancer-Associated SF3B1 Hotspot Mutations Induce Cryptic 3′ Splice Site Selection through Use of a Different Branch Point. Cell Reports, 13, 1033–1045.

57. Shiozawa, Y., Malcovati, L., Gallì, A., Sato-Otsubo, A., Kataoka, K., Sato, Y., Watatani, Y., Suzuki, H., Yoshizato, T., Yoshida, K., et al. (2018) Aberrant splicing and defective mRNA production induced by somatic spliceosome mutations in myelodysplasia. Nat Commun, 9, 3649.

58. Alsafadi, S., Houy, A., Battistella, A., Popova, T., Wassef, M., Henry, E., Tirode, F., Constantinou, A., Piperno-Neumann, S., Roman-Roman, S., et al. (2016) Cancer-associated SF3B1 mutations affect alternative splicing by promoting alternative branchpoint usage. Nat Commun, 7, 10615.

59. Kesarwani, A.K., Ramirez, O., Gupta, A.K., Yang, X., Murthy, T., Minella, A.C. and Pillai, M.M. (2017) Cancer-associated SF3B1 mutants recognize otherwise inaccessible cryptic 3′ splice sites within RNA secondary structures. Oncogene, 36, 1123–1133.

60. Furney, S.J., Pedersen, M., Gentien, D., Dumont, A.G., Rapinat, A., Desjardins, L., Turajlic, S., Piperno-Neumann, S., De La Grange, P., Roman-Roman, S., et al. (2013) *SF3B1* Mutations Are Associated with Alternative Splicing in Uveal Melanoma. Cancer Discovery, 3, 1122–1129.

61. Wang, L., Brooks, A.N., Fan, J., Wan, Y., Gambe, R., Li, S., Hergert, S., Yin, S., Freeman, S.S., Levin, J.Z., et al. (2016) Transcriptomic Characterization of SF3B1 Mutation Reveals Its Pleiotropic Effects in Chronic Lymphocytic Leukemia. Cancer Cell, 30, 750–763.

62. Zhang, J., Lieu, Y.K., Ali, A.M., Penson, A., Reggio, K.S., Rabadan, R., Raza, A., Mukherjee, S. and Manley, J.L. (2015) Disease-associated mutation in *SRSF2* misregulates splicing by altering RNA-binding affinities. Proc. Natl. Acad. Sci. U.S.A., 112.

63. Kim, E., Ilagan, J.O., Liang, Y., Daubner, G.M., Lee, S.C.-W., Ramakrishnan, A., Li, Y., Chung, Y.R., Micol, J.-B., Murphy, M.E., et al. (2015) SRSF2 Mutations Contribute to Myelodysplasia by Mutant-Specific Effects on Exon Recognition. Cancer Cell, 27, 617– 630.

64. Liang, Y., Tebaldi, T., Rejeski, K., Joshi, P., Stefani, G., Taylor, A., Song, Y., Vasic, R., Maziarz, J., Balasubramanian, K., et al. (2018) SRSF2 mutations drive oncogenesis by activating a global program of aberrant alternative splicing in hematopoietic cells. Leukemia, 32, 2659–2671.

65. Okeyo-Owuor, T., White, B.S., Chatrikhi, R., Mohan, D.R., Kim, S., Griffith, M., Ding, L., Ketkar-Kulkarni, S., Hundal, J., Laird, K.M., et al. (2015) U2AF1 mutations alter sequence specificity of pre-mRNA binding and splicing. Leukemia, 29, 909–917.

66. Ilagan, J.O., Ramakrishnan, A., Hayes, B., Murphy, M.E., Zebari, A.S., Bradley, P. and Bradley, R.K. (2015) *U2AF1* mutations alter splice site recognition in hematological malignancies. Genome Res., 25, 14–26.

67. Graubert, T.A., Shen, D., Ding, L., Okeyo-Owuor, T., Lunn, C.L., Shao, J., Krysiak, K., Harris, C.C., Koboldt, D.C., Larson, D.E., et al. (2012) Recurrent mutations in the U2AF1 splicing factor in myelodysplastic syndromes. Nat Genet, 44, 53–57.

68. Brooks, A.N., Choi, P.S., De Waal, L., Sharifnia, T., Imielinski, M., Saksena, G., Pedamallu, C.S., Sivachenko, A., Rosenberg, M., Chmielecki, J., et al. (2014) A Pan-Cancer Analysis of Transcriptome Changes Associated with Somatic Mutations in U2AF1 Reveals Commonly Altered Splicing Events. PLoS ONE, 9, e87361.

69. Chi, Z., Gupta, V. and Query, C. (2025) U2-2 snRNA Mutations Alter the Transcriptome. 10.1101/2025.03.12.642709.

70. Suzuki, H., Kumar, S.A., Shuai, S., Diaz-Navarro, A., Gutierrez-Fernandez, A., De Antonellis, P., Cavalli, F.M.G., Juraschka, K., Farooq, H., Shibahara, I., et al. (2019) Recurrent noncoding U1 snRNA mutations drive cryptic splicing in SHH medulloblastoma. Nature, 574, 707– 711.

71. Shuai, S., Suzuki, H., Diaz-Navarro, A., Nadeu, F., Kumar, S.A., Gutierrez-Fernandez, A., Delgado, J., Pinyol, M., López-Otín, C., Puente, X.S., et al. (2019) The U1 spliceosomal RNA is recurrently mutated in multiple cancers. Nature, 574, 712–716.

