## Supplementary Figures and Legends for "The human branchpoint-interacting stem loop sequence and structure regulates U2 snRNA expression, branchpoint recognition, and transcriptome"

Supplementary Figure 1

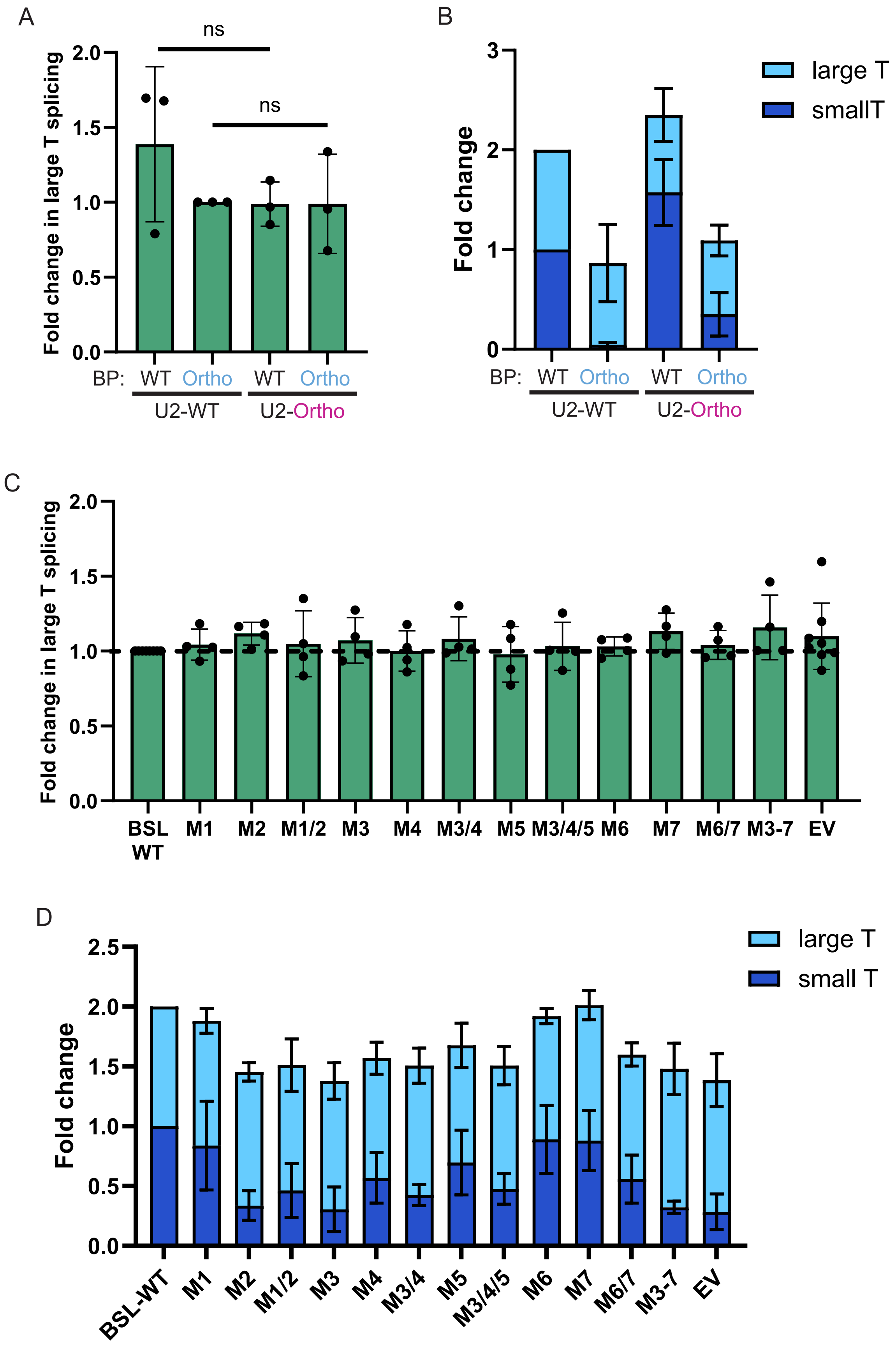

Supplementary Figure 2

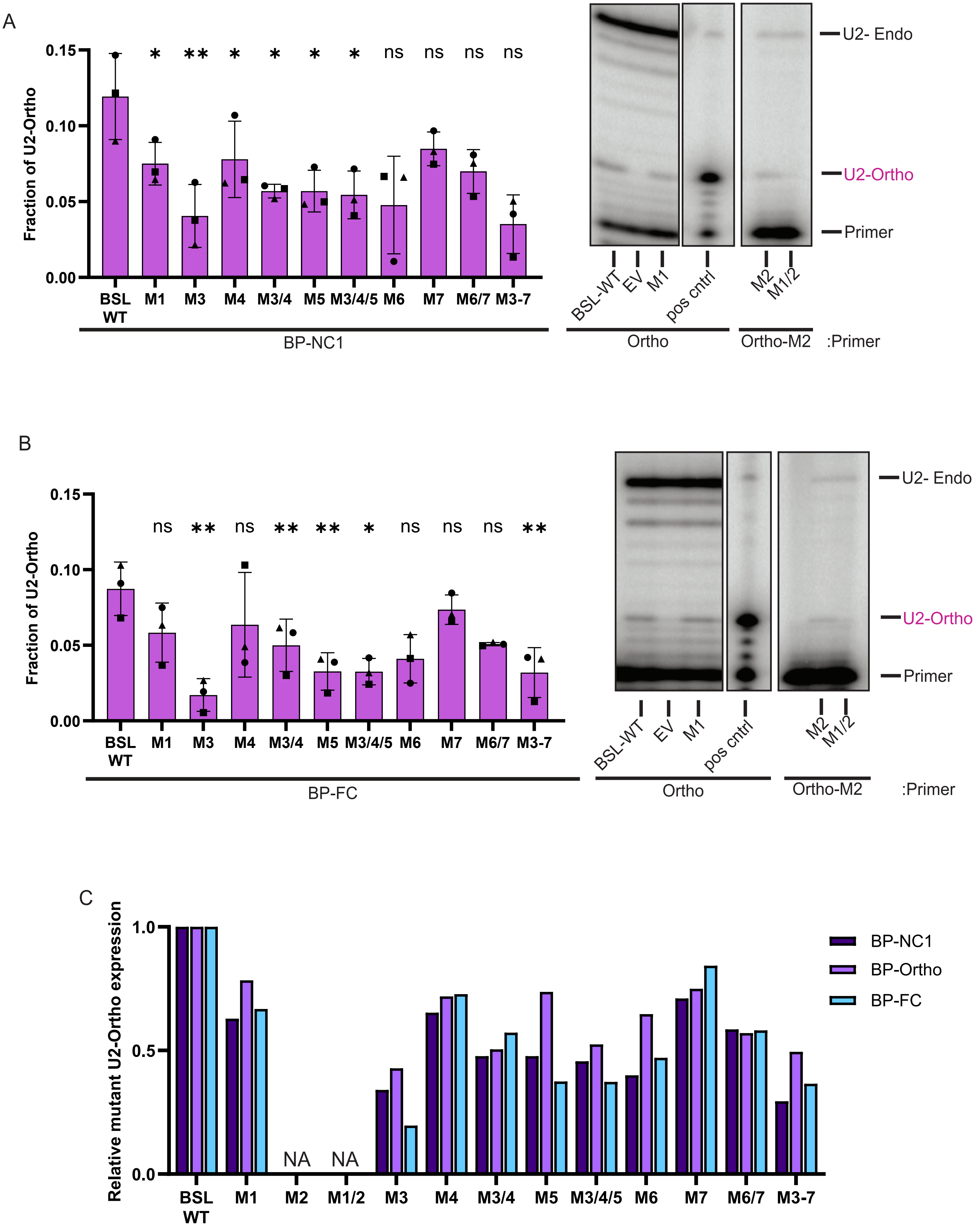

Supplementary Figure 3

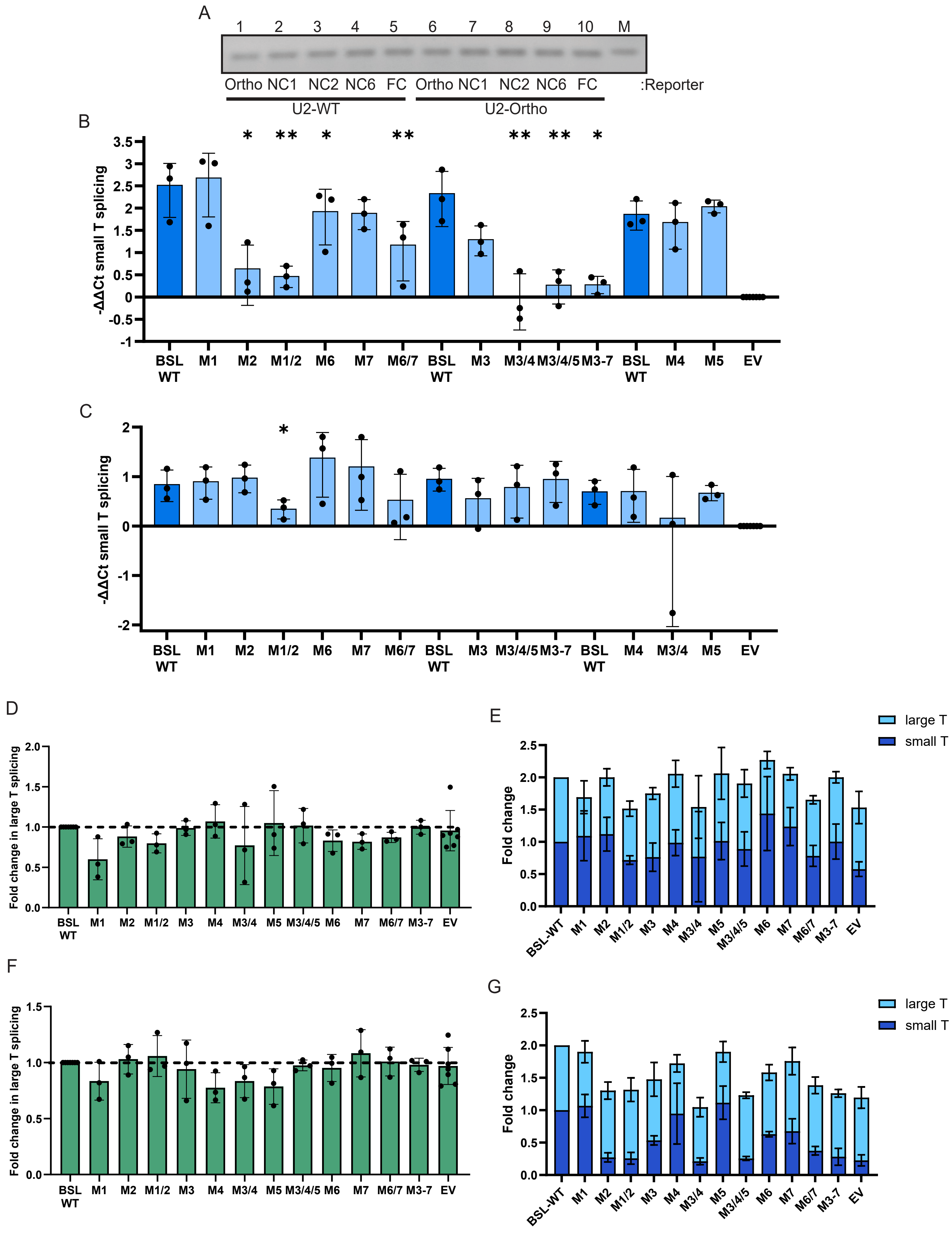

### Supplementary Figure 4

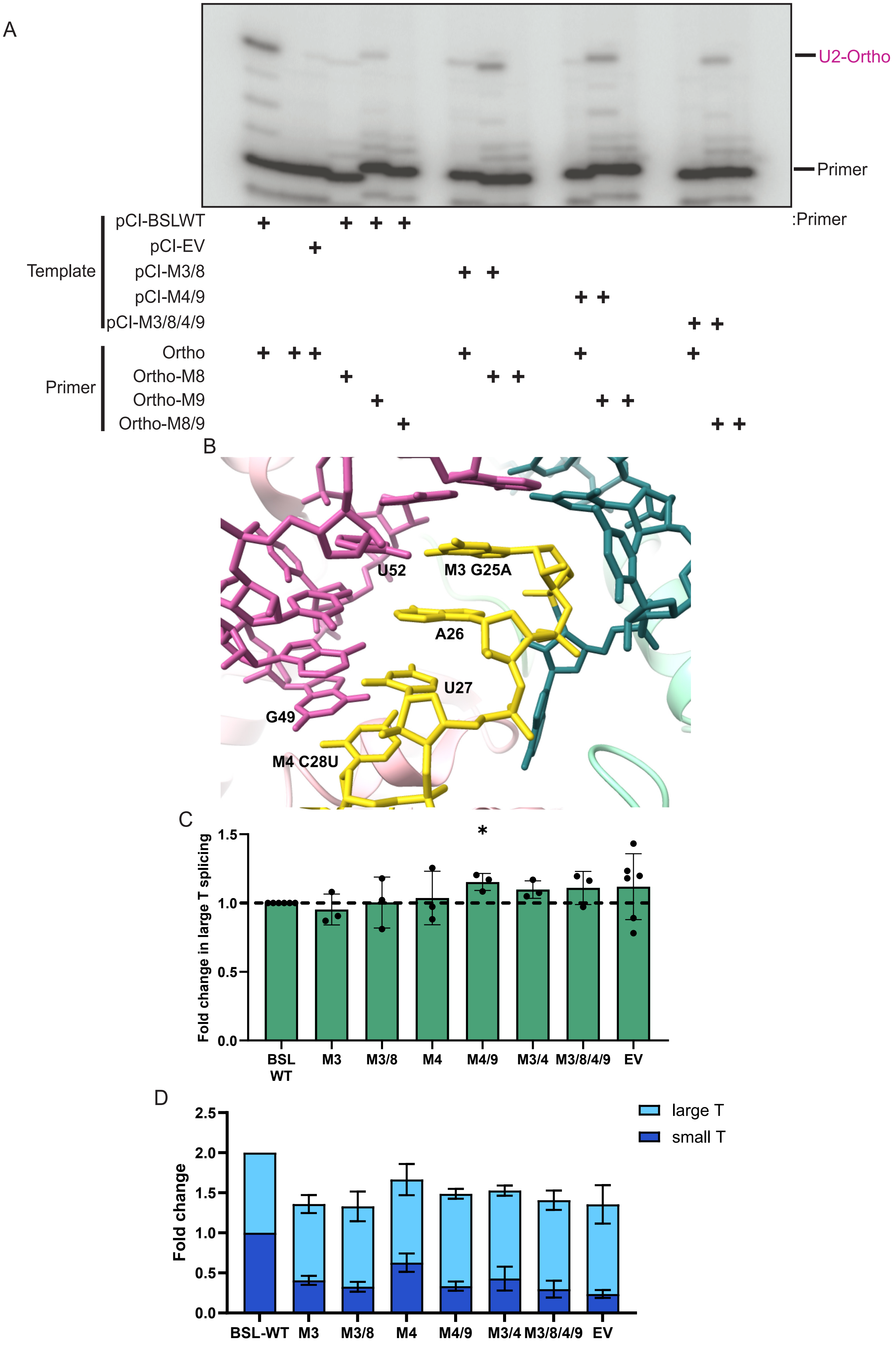

Supplementary Figure 5

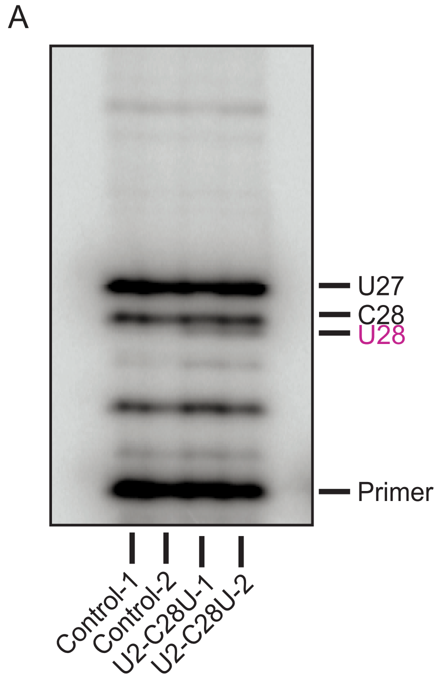


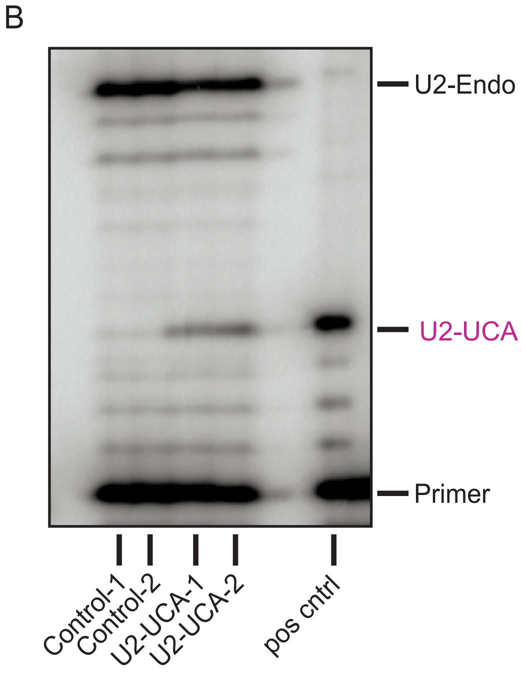

#### SUPPLEMENTARY FIGURE LEGENDS

**Supplementary Figure 1.** Perturbing BSL base-pairing potential decreases U2 snRNA stable expression and reporter splicing. (**A**) Difference in large T intron splicing by RT-qPCR for samples shown in Figure 1C and D. -ΔΔCt is derived from large T splice junction probe normalized to exon2 and relative to sample 2 (BP-Ortho and U2-WT); n=3. (**B**) RT-qPCR fold change normalized ratios of large and small T splicing of cell transfections of Figure 1C and D. Lanes are normalized to sample 1 (BP-WT and U2-WT); n=3. (**C**) Difference in large T intron splicing by RT-qPCR for samples shown in Figure 1F, G, H, and I. -ΔΔCt is derived from large T splice junction probe normalized to exon2 and relative to BSL-WT; n=4. (**D**) RT-qPCR fold change normalized ratios of large and small T splicing of cell transfections of Figure 1F, G, H and I. Lanes are normalized to BSL-WT of the same transfection batch; n=4. In all cases, error bars represent standard deviation.

**Supplementary Figure 2.** U2 snRNA expression remains unchanged across branchpoint reporter transfections. (**A**) Expression of orthogonal U2 snRNA BSL WT and mutations after transfection with branchpoint reporter BP-NC1 (Figure 2D and E). Left: Fraction of U2-Ortho relative to total U2 snRNA quantified from primer extension stops. Right: Representative PAGE of radiolabeled U2 snRNA primer extension analysis from cells transfected with BSL-WT and stabilizing mutants. Extension stops for endogenous/wildtype U2 snRNA and U2-Ortho are labeled; n=3. (**B**) Expression of orthogonal U2 snRNA BSL WT and mutations after transfected with branchpoint reporter BP-FC (Figure 2D and E). Left: Fraction of U2-Ortho relative to total U2 snRNA quantified from primer extension stops. Right: Representative PAGE of radiolabeled U2 snRNA primer extension analysis from cells transfected with BSL-WT and stabilizing mutants. Extension stops for endogenous/wildtype U2 snRNA and U2-Ortho are labeled; n=3. (**C**) Bar graph of relative orthogonal BSL mutant U2 snRNA expression from transfection of U2 snRNA BSL-WT or BSL mutant with each orthogonal small T branchpoint reporter (Figure 1F, G I and H, Figure 2D and E, and (A) and (B)). Lanes are relative to BSL-WT cells of the same transfection. Expression of mutations M2 and M1/2 could not be accurately quantified by primer extension. In all cases, error bars represent standard deviation.

**Supplementary Figure 3.** BSL base-pairing potential influences branchpoint sequence recognition (**A**) Representative agarose gel of large T RT-PCR from transfection in Figure 2B and C. DNA ladder band is indicated by M. (**B**) Difference in small T intron splicing by RT-qPCR for samples transfected with BP-FC shown in Figure 2D and E, Supplementary Figure 2B and C. -ΔΔCt is derived from small T splice junction probe normalized to exon2 and relative to empty vector (EV). Lanes are grouped by transfection batch for comparison.; n=3. (**C**) Difference in small T intron splicing by RT-qPCR for samples transfected with BP-NC1 shown in Figure 2D and E, Supplementary Figure 2A and C. -ΔΔCt is derived from small T splice junction probe normalized to exon2 and relative to empty vector (EV). Lanes are grouped by transfection batch for comparison.; n=3. (**D**) Difference in large T intron splicing by RT-qPCR for samples transfected with BP-NC1 shown in Figure 2D and E and Supplementary Figure 2A and C and (C). -ΔΔCt is derived from large T splice junction probe normalized to exon2 and relative to BSL-WT; n=3. (**E**) RT-qPCR fold change normalized ratios of large and small T splicing of cell transfections with BP-NC1 shown in Figure 2D and E and Supplementary Figure 2A and C and (C). Lanes are normalized to BSL-WT of the same transfection batch; n=3. (**F**) Difference in large T intron splicing by RT-qPCR for samples transfected with BP-FC shown in Figure 2D and E and Supplementary Figure 2B and C and (B). -ΔΔCt is derived from large T splice junction probe normalized to exon2 and relative to BSL-WT; n=3. (**G**) RT-qPCR fold change normalized ratios of large and small T splicing of cell transfections with BP-FC shown in Figure 2D and E and Supplementary Figure 2B and C and (B). Lanes are normalized to BSL-WT of the same transfection batch; n=3. In all cases, error bars represent standard deviation.

**Supplementary Figure 4.** Compensatory mutations do not reverse the negative effect of some BSL weakening mutations. (**A**) Representative PAGE of radiolabeled U2 snRNA primer extension analysis for the indicated BSL mutants. Extension stops for endogenous/wildtype U2 snRNA and U2-Ortho are labeled. (**B**) Model of U2-U6 snRNA Helix Ia from Complex Bact (PDB 6FF4) show the position of M3 and M4 mutations. U2 snRNA is in yellow and U6 snRNA is in magenta. (**C**) Difference in large T intron splicing by RT-qPCR for samples shown in Figure 3B and C. -ΔΔCt is derived from large T splice junction probe normalized to exon2 and relative to BSL-WT; n=3. (**D**) RT-qPCR fold change normalized ratios of large and small T splicing of cell transfections of Figure 3B and C and (C). Lanes are normalized to BSL-WT of the same transfection batch; n=3. In all cases, error bars represent standard deviation.

**Supplementary Figure 5:** Expression of U2 snRNA BSL mutations results in global alterations to splicing and gene expression. (**A**) Representative PAGE of radiolabeled U2 snRNA primer extension analysis from HEK293T cells transduced with empty vector (Control) and U2 snRNA C28U (U2-C28U). Extension stops for endogenous/wildtype U2 snRNA and U2-C28U are labeled. (**B**) Representative PAGE of radiolabeled U2 snRNA primer extension analysis from HEK293T cells transduced with empty vector (Control) and U2-Ortho (U2-UCA). Extension stops for endogenous/wildtype U2 snRNA and U2-UCA are labeled.

**Supplementary Table Legends**

**Supplementary Table 1.** Primer sequences used in this study,

**Supplementary Table 2.** Control versus U2-C28U Differential Splicing Events. junctionCounts DEXSeq output of significantly differential splicing events. gene : gene symbol, event_id : unique name for event, event_type : type of event, chr : chromosome, start : start coordinate of event, end : end coordinate of event, strand : strand, cntl_mean_ijc : mean read count of all junctions in the included form across replicates in control cells, cntl_mean_ejc : mean read count of all junctions in the excluded (skipped) form across replicates in control cells, cntl_mean_psi : mean of PSI values computed for all pairs of included, excluded junction counts across replicates in control cells, cmut_mean_ijc : mean read count of all junctions in the included form across replicates in U2-C28U cells, cmut_mean_ejc : mean read count of all junctions in the excluded (skipped) form across replicates in U2-C28U cells, cmut_mean_psi : mean of PSI values computed for all pairs of included, excluded junction counts across replicates in U2-C28U cells, dpsi : difference of mean(PSI) between conditions, event_qval : Q-value describing the positive false discovery rate of the difference of mean(PSI) between conditions, sig : 1 = event is significantly more included (dpsi ≥ 0.1, event_qval ≤ 0.05) in U2-C28U cells relative to control cells. -1 = event is significantly less included (dpsi ≤ -0.1, event_qval ≤ 0.05) in U2-C28U cells relative to control cells.

**Supplementary Table 3.** Control versus U2-UCA Differential Splicing Events. junctionCounts DEXSeq output of significantly differential splicing events. gene : gene symbol, event_id : unique name for event, event_type : type of event, chr : chromosome, start : start coordinate of event, end : end coordinate of event, strand : strand, cntl_mean_ijc : mean read count of all junctions in the included form across replicates in control cells, cntl_mean_ejc : mean read count of all junctions in the excluded (skipped) form across replicates in control cells, cntl_mean_psi : mean of PSI values computed for all pairs of included, excluded junction counts across replicates in control cells, acu_mean_ijc : mean read count of all junctions in the included form across replicates in U2-UCA cells, acu_mean_ejc : mean read count of all junctions in the excluded (skipped) form across replicates in U2-UCA cells, acu_mean_psi : mean of PSI values computed for all pairs of included, excluded junction counts across replicates in U2-UCA cells, dpsi : difference of mean(PSI) between conditions, event_qval : Q-value describing the positive false discovery rate of the difference of mean(PSI) between conditions, sig : 1 = event is significantly more included (dpsi ≥ 0.1, event_qval ≤ 0.05) in U2-UCA cells relative to control cells. -1 = event is significantly less included (dpsi ≤ -0.1, event_qval ≤ 0.05) in U2-UCA cells relative to control cells.

**Supplementary Table 4.** Control versus U2-C28U Differentially Upregulated Genes. DESeq2 output of significantly upregulated genes. ENSG: Ensembl gene ID, GeneName: Gene Symbol, baseMean: normalized count value average, log2FoldChange: log2 of fold change between C28U and control cells, lfcSE: standard error estimate, stat: test statistic value, pvalue: P-value statistic, padj: Adjusted P-value statistic.

**Supplementary Table 5.** Control versus U2-UCA Differentially Upregulated Genes. DESeq2 output of significantly upregulated genes. ENSG: Ensembl gene ID, GeneName: Gene Symbol, baseMean: normalized count value average, log2FoldChange: log2 of fold change between C28U and control cells, lfcSE: standard error estimate, stat: test statistic value, pvalue: P-value statistic, padj: Adjusted P-value statistic.
